# Gating transitions and modulation of a hetero-octameric AMPA glutamate receptor

**DOI:** 10.1101/2020.12.13.422579

**Authors:** Danyang Zhang, Jake F. Watson, Peter M. Matthews, Ondrej Cais, Ingo H. Greger

## Abstract

AMPA glutamate receptors (AMPARs) mediate the majority of excitatory transmission in the brain, and enable synaptic plasticity that underlies learning ^1^. A diverse array of AMPAR signaling complexes are established by receptor auxiliary subunits, associating in various combinations to modulate trafficking, gating and synaptic strength ^2^. However, their mechanisms of action are poorly understood. Here, we determine cryo-electron microscopy structures of the heteromeric GluA1/2 receptor assembled with both TARP-γ8 and CNIH2, the predominant AMPAR complex in the forebrain, in both resting and active states (at 3.2 and 3.7 Å, respectively). Consequential for gating regulation, two γ8 and two CNIH2 subunits lodge at distinct sites beneath the ligand-binding domains of the receptor tetramer, with site-specific lipids shaping each interaction. Activation leads to a stark asymmetry between GluA1 and GluA2 along the ion conduction path, and an outward expansion of the channel triggers counter-rotations of both auxiliary subunit pairs, that promotes the active-state conformation. In addition, both γ8 and CNIH2 pivot towards the pore exit on activation, extending their reach for cytoplasmic receptor elements. CNIH2 achieves this through its uniquely extended M2 helix, which has transformed this ER-export factor into a powerful positive AMPAR modulator, capable of providing hippocampal pyramidal neurons with their integrative synaptic properties.

## Introduction

Unique amongst ionotropic glutamate receptors, AMPARs form a diverse array of signaling complexes, specialized for a range of functions - from faithfully decoding high frequency inputs to integration of low-frequency signals in support of synaptic plasticity ^3^. This spectrum results from a mosaic of receptor combinations, assembled from four pore-forming subunits (GluA1-4) and multiple auxiliary proteins that exist in various stoichiometries and exhibit distinct expression patterns in the brain ^4,5^. The predominant subunit, GluA2, dictates critical functions such as ion permeation, and forms heteromeric complexes with GluA1 or GluA3 in principal neurons of the forebrain ^2,6^.

Members of the transmembrane AMPAR regulatory proteins (TARPs γ2, 3, 4, 5, 7, 8) and cornichon homologue (CNIH2, 3) families are the most widely expressed auxiliary subunits ^2^. TARP γ8 and CNIH2 are highly abundant at cortical synapses ^7,8^, and are powerful AMPAR modulators ^9,10^ that co-purify and synergize to determine receptor expression levels, kinetics and pharmacology ^7,11–13^. Both are four-helical bundle transmembrane proteins, yet they have different topologies, resulting in fundamentally different mechanisms of action. Whilst engaging the receptor transmembrane domain (TMD), TARPs possess an elaborate extracellular beta-sheet structure that contacts the ligand-binding domains (LBDs) ^14–16^. CNIHs on the other hand are mostly embedded within the membrane ^17^, but have two cytoplasmic loop elements of unknown structure that contribute to their function as cargo exporters from the endoplasmic reticulum (ER) ^18^. Hetero-auxiliary receptor complexes have yet to be resolved, and therefore the stoichiometry and distinct modulatory mechanism of each associating protein are poorly understood. To address these questions, we investigated the architecture and functioning of the principal forebrain AMPAR complex, the GluA1/2 receptor assembled with TARP γ8 and CNIH2.

### Recombinant GluA1/2/γ8/CNIH2 resembles native AMPARs

We first reconstituted the hetero-auxiliary AMPAR complex and compared its properties to those of native hippocampal AMPARs. CNIH2-expressing HEK-Expi293F cells were co-transfected with GluA1 and a GluA2_γ8 (A2_γ8) fusion gene, to ensure the presence of two γ8 subunits ^15^. Although AMPAR stoichiometries with four γ8 molecules may exist ^19^, the co-existence of γ8 and CNIH2 predominates in forebrain neurons ^7,11^. In response to rapid glutamate application to excised membrane patches, γ8 slowed desensitization rates about 2-fold compared to GluA1/2 alone, and addition of CNIH2 led to much further slowing that was greater than that of four γ8 subunits alone (**Fig. 1a, b**) ^15^. CNIH2 also further increased the equilibrium current (**Extended Data Fig. 1a**). Together with previous data ^13^, this confirms incorporation of CNIH2 into a hetero-auxiliary complex.

**Figure. 1:**
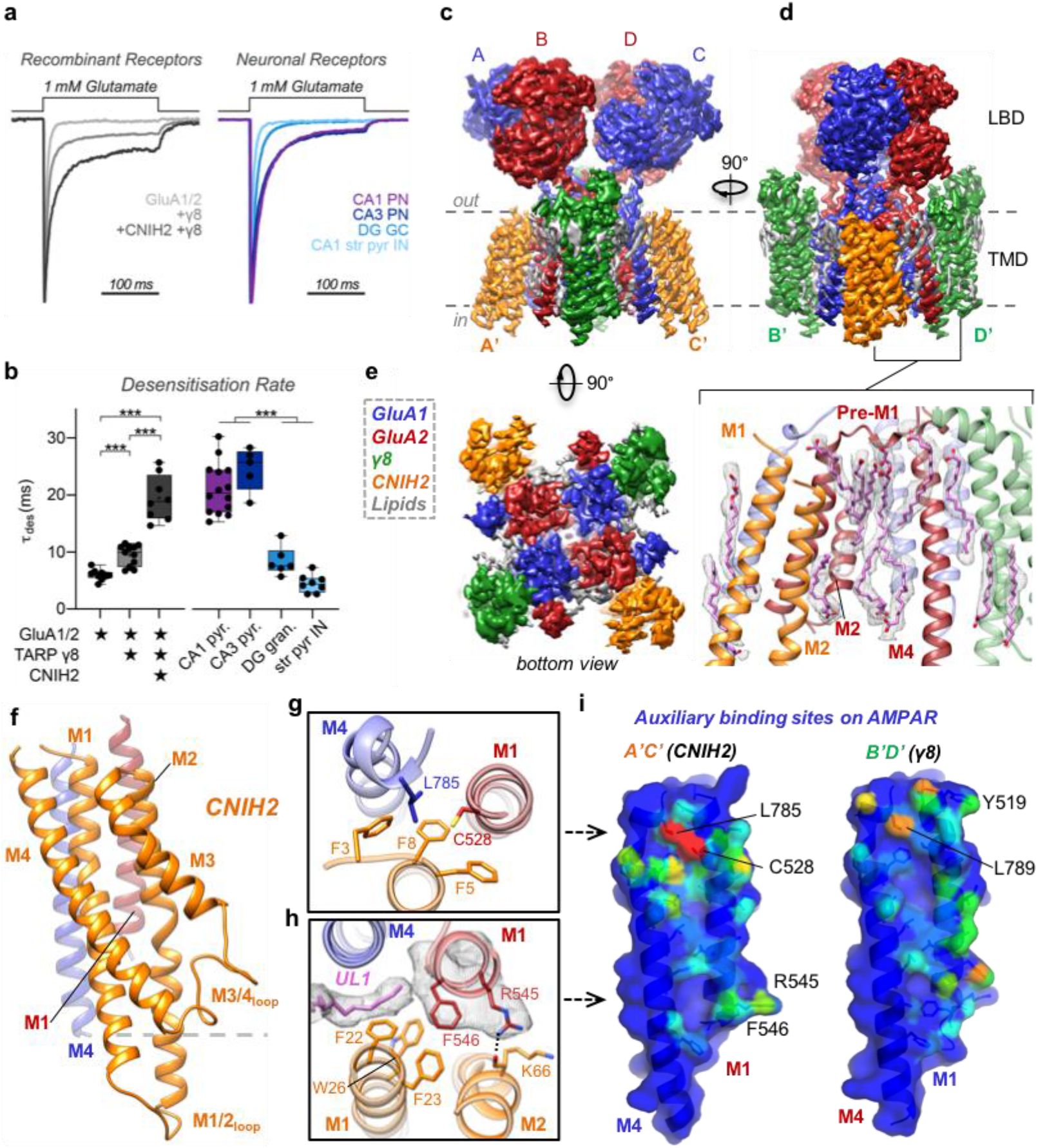
Physiology and architecture of the GluA1/2_γ8 /CNIH2 complex. **a,b**, Auxiliary subunits slow desensitization rates of recombinant receptors (**a**, left). Native AMPAR responses show diverse properties (**a**, right), with A1/2_γ8 +CNIH2 recapitulating CA1 & CA3-like kinetics (**b**, weighted τ_des_ (ms), mean ± SEM). *Recombinant receptors:* GluA1/2: 6.01 ± 0.31, n=9; +γ8: 9.29 ± 0.53, n=12; +CNIH2+γ8:19.71 ± 1.27, n=9. *Neuronal receptors:* CA1 pyramidal: 21.03 ± 1.19, n=14; CA3 pyramidal 24.51 ± 1.69, n=5; DG granule cell: 8.38 ± 1.01, n=6; CA1 stratum pyramidale interneurons: 4.48 ± 0.55, n=8; Welch’s ANOVA tests with Dunnett’s multiple comparisons test – Recombinant: W(2,15.11) = 60.68, p<0.0001; Neurons: W(3,11.60) = 76.28, p<0.0001). **c-e**, Cryo-EM maps of the AMPAR octamer, depicting the LBD and TMD domain layers. Core subunits positioned to AC/BD, and auxiliary subunits associated with A’C’/B’D’ sites are shown. (**c**) Front view, depicting γ8 located at B’D’ sites. (**d**) Side view, offering a view of CNIH2 at A’C’ sites, beneath an LBD dimer. Inset: annular lipids concentrating at side of the TMD, beneath the pre-M1 helix. (**e**) Bottom view, highlighting CNIH2 binding to GluA2 transmembrane helices M1-3 (red), and γ8 contacting the GluA1 M1-3 helices (blue). **f**, Model of CNIH2, including the M1/2 and M3/4 cytosolic loops, docking to its binding site (M1_GluA2_ [red] M4_GluA1_ [blue]). G, CNIH2 Phe3, −5, −8 slotting into the top tier of its binding site close to Cys528 (GluA2) and Leu785 (GluA1). **h**, CNIH2 contacts at the bottom of the binding site are mediated by Phe23 and Lys66; a lipid (UL1) penetrates the A’C’ site and interacts with Phe546 (GluA2). **i**, A’C’ (left) and B’D’ (right) site surface representation. M1_A2_ and M4_A1_ residues contacted by CNIH2 are coloured depending on the number of atoms contributing to the interaction (red: high, blue: low). Contacts were counted using the ‘findNeighbors’ command from *ProDy* ^38^, with a cutoff of 4.5 Å between heavy atoms.

We found close correspondence between responses from the recombinant complex and AMPARs expressed in CA1 and CA3 pyramidal neurons, but not to those from dentate granule cells or CA1 interneurons, where γ8 and other auxiliary subunits prevail (**Fig. 1a, b**, **Extended Data Fig. 1a**) ^20^. This demonstrates the diversity of AMPAR complexes, and provides further evidence that CNIH2 acts to define the unique response properties of pyramidal neurons, with the GluA1/2/γ8/CNIH2 octamer the major assortment in these cells ^7,11,13^. We subjected this native-like complex for structural analysis by cryo-electron microscopy (cryo-EM).

### Trapping GluA1/2/γ8/CNIH2 in active and inactive states

To resolve gating transitions, the complex was trapped both in an inactive state, bound to the competitive antagonist NBQX, and in an active state with the agonist L-glutamate together with the desensitization blocker cyclothiazide (CTZ) (**Extended Data Fig. 1b**). 3-D reconstructions revealed the classic three-layered AMPAR domain architecture, with the extracellular N-terminal domain (NTD) and LBD arranged as dimers of dimers, attached to the ion channel of pseudo four-fold symmetry (**Fig. 1c-e, Extended Data Fig. 1c**). Focused processing of the LBD-TMD, and of individual domain layers resulted in high-resolution maps ranging from 3.0 Å to 3.8 Å (**Extended Data Fig. 2-4**, and **Extended Data Table 1**), permitting us to follow gating transitions at comparable resolutions. Moreover, reconstruction of particles from resting-state data lacking CNIH2, enabled us to also examine the effects of CNIH2-association with the receptor (**Extended Data Fig. 2**).

### Subunit arrangement in the hetero-octamer

EM-map quality permitted unambiguous assignment of the four core subunits ^15,21^, placing GluA1, marked by unique N-glycans in the NTD (**Extended Data Fig. 5a**), to the two outer (or AC) positions, and GluA2 to the inner (BD) positions. This arrangement dictates function as the BD subunits exert a greater mechanical force on the ion channel gate than the AC subunits ^22^, with consequences on the entire complex, as we describe below. The core subunits alternate around the pore with their peripheral M1 and M4 transmembrane helices contributing two pairs of non-equivalent binding sites for auxiliary subunits: the A’C’ sites (M1_GluA2_/M4_GluA1_) locate beneath the LBD dimers, thus these positions are spatially more restricted than the B’D’ sites (M1_GluA1_/M4_GluA2_), beneath the LBD dimer-of dimers interface (**Fig. 1c-e**). With γ8 binding to its preferential B’D’ site ^15^, CNIH2 occupies the A’C’ positions, and despite its greater detergent resistance ^4,7^, CNIH2 does not displace γ8 from the receptor. This organization determines gating modulation, as each site provides differential access of the auxiliaries to the gating machinery.

We obtained a complete structural model for CNIH2, including its two sequence-diverse cytoplasmic loops between the M1/2 and the M3/4 helices, that were not resolved in the recent CNIH3 structure ^17^ (**Extended Data Fig. 5f**). Lodging to the A’C’ sites, the CNIH2 M1 and M2 helices project deeply into the cytoplasm (>20 Å) (**Fig. 1f**), forming a loop element of currently unknown function. A comparison of the binding modes for CNIH2 and γ8 shows that both engage the same set of residues on the receptor, yet their footprints vary and differences in side chain orientations along the M1 helix are apparent (**Fig. 1g-i, Extended Data Fig. 6a**). While TARP γ8 contacts are spread out more evenly, CNIH2 prominently engages two regions: the upper tier of the A’C’ site, centered around GluA2 Cys528, and its base adjacent to GluA2 Phe546 (**Fig. 1i**). Three highly conserved N-terminal phenylalanines (Phe3, −5, −8)^17^ slot CNIH2 into the TMD proximal to the gate (**Fig. 1g**), and these are crucial for modulation, as we show below. At the base, CNIH2 interacts with Arg545 and Phe546 through polar and hydrophobic contacts (**Fig. 1h**), and this region of the A’C’ site is shaped by lipids.

### Annular lipids shape auxiliary subunit binding sites

An abundance of lipid-like densities ensheath the ion channel, forming an annular belt demarcating the membrane boundaries (inset in **Fig. 1d**). Lipids in the upper leaflet connect the gate-surrounding pre-M1 helices to the M2 pore helices in the lower leaflet, their location within the lateral fenestrations of the channel indicate a role in channel modulation. Other lipids, denoted lower-leaflet lipid (LL) 1 and LL2, bridge between auxiliary subunits and the M2 pore helix, thereby extending the reach of these modulating proteins to the ion conduction path (**Extended Data Fig. 6c, d**).

The acyl chains of two lipids penetrate the lower part of the CNIH2 binding site: wedging between GluA1 and CNIH2, upper-leaflet lipid (UL) 1 contacts Phe546 (**Fig. 1h**), while immediately beneath, LL3 anchors CNIH2 via its Phe22 and Trp26 side chains to the base of the GluA1 M4 helix (**Fig. 1i, Extended Data Fig. 6d**). These acyl chains likely influence CNIH2 binding by extending the hydrophobic network at the A’C’ site. This idea is supported by the GluA2-CNIH3 structure (PDB 6PEQ), where some lipids are also observed ^17^, but LL1 and LL2 are missing. As a result, CNIH3 aligns with the receptor more closely than CNIH2 (**Extended Data Fig. 6e**). Hence, lipids may regulate CNIH-modulation of AMPARs.

### The GluA2 subunit dominates gating

Receptor activation is triggered by glutamate closing the LBD ‘clamshells’ ^23^ (**Extended Data Fig. 5b-e**), leading to separation of the lower lobes within an LBD heterodimer and rearrangement of the LBD-TMD linkers to open the gate ^14,16^. This state transition is well resolved. Focusing on the outer gate formed by the M3 transmembrane helices, M3 linker tension ruptures polar contacts between GluA1 Arg624 and GluA2 Arg628, a position determining gating kinetics ^24^, and breaks a stacking interaction between GluA1 Arg624 and Phe623 in GluA2 (**Fig. 2a, b**). These changes are stabilized by reorganization of the gating linkers, and are accompanied by opening of the channel gate formed by hydrophobic contacts facing the pore axis along the conserved SY**<u>T</u>** ANL**<u>A</u>** AFL**<u>T</u>** motif (**Fig. 2b, c**).

**Figure. 2:**
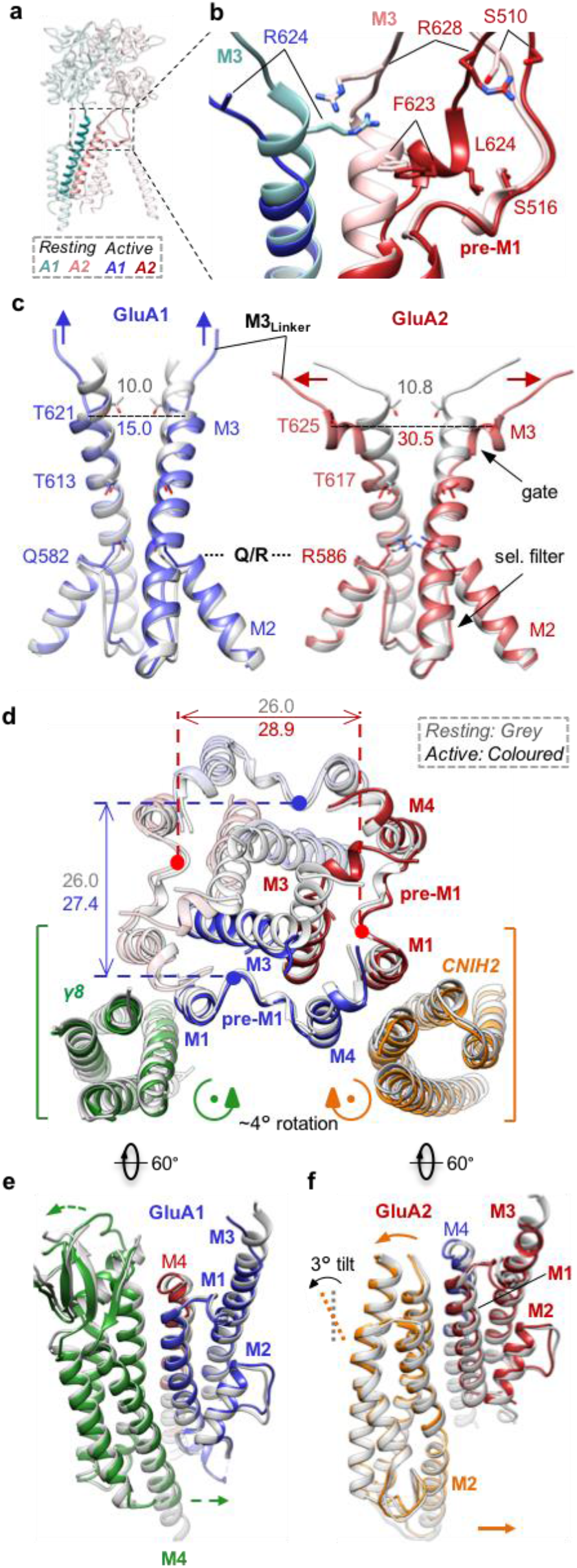
Gating transitions of the A1/2_γ8/C2 octamer. **a**, View onto a GluA1/GluA2 LBD dimer and associated M3 gating helices (depicted in bold) in the resting state. **b**, Side view onto the M3 gate with open (bold colour) and resting (soft colour) states overlain, depicting rearrangement of the M3 linkers and reorientation of M3 side chins relative to pre-M1. **c**, Comparison of GluA1 and GluA2 conduction path; resting state (grey) is superimposed on the active state (colour). Divergence from the closed gate is apparent at the top of M3, and is more pronounced in GluA2 (Cα distances between opposing T621 (GluA1), and T625 (GluA2) indicated). Also shown is the selectivity filter with the Q/R site at its apex. **d**, Top view onto the closed-(grey) and open-state (colour) models at the level of the M3 gate. The dilation of the gate-surrounding helices is shown and measured between opposing pre-M1 helices. **e**, Side view showing outward expansion of the upper part of GluA1 M1 and M3 and the γ8 β-sheet, accompanied by a slight inward movement of the γ8 M4 helix towards the GluA1 M12 loop. **f**, CNIH2 undergoes a ~3° pivot (‘tilt’) on activation, moving its M12 loop closer to the pore axis.

Gate dilation is more pronounced between the two opposing GluA2 subunits than between GluA1, and this continues into the selectivity filter, formed by the M2 pore loops midway in the conduction path (**Fig. 2c**). At the Q/R site (Arg586), which determines Ca^2+^ flux and channel conductance at the apex of the M2 pore loop ^25,26^, activation widens the filter entrance selectively between the GluA2 subunits (**Extended Data Fig. 7a, b**). This asymmetric widening extends down to GluA2 Cys589, midway in the filter and to the M2 pore helices, which separate by > 1 Å in GluA2 but not GluA1, and is apparent at the cytoplasmic face of the pore, the site modulated by polyamines ^27^ (**Extended Data Fig. 7c, d, e**). We note that binding of CNIH2, which increases channel conductance ^28^, does not cause a visible expansion of the selectivity filter under resting-state conditions (**Extended Data Fig. 7b-d**).

Receptor activation is also accompanied by an anti-clockwise twist of the AMPAR TMD helices, when looking down the pore from the extracellular side, and a marked expansion in the upper part of the channel that surrounds the gate (**Extended Data Fig. 8a)**. This dilation is greater between opposing GluA2 subunits, than between GluA1, and includes the channel’s peripheral helices (pre-M1, M1 and M4) and both pairs of auxiliary subunits (**Fig. 2d**).

### Gating transitions of core and auxiliary subunits

Dictated by its architecture, receptor activation differentially impacts the interactions between core and auxiliary subunits. On activation, the GluA2 M3 gate helix pushes its Phe623 and Leu624 side chains against its pre-M1 helix, triggering channel dilation (**Fig. 2b, d, Supplementary Video 1**). Pre-M1 couples LBD closure to channel opening, and shapes gating kinetics throughout iGluRs ^29–31^. At the same time, the GluA2 M1 and M3 linkers closely align and approach the TARP extracellular (β4) loop, which is simultaneously contacted by the GluA2 LBD through its ‘KGK’ motif ^32^ (**Extended Data Fig. 8b**). GluA1 does not experience these large changes, as its M3 linkers are pulled upward (left panel in **Fig. 2c**), a characteristic of AC subunits ^14,16^. As a result, the GluA1 M3 helices minimally contact pre-M1 on activation, which underlies the asymmetric expansion of the channel (**Fig. 2d**). We also observe newly formed contacts between the GluA1 M4 linkers (at Thr780) with the GluA2 M3 gate on activation, a change that is not apparent with GluA1 M3. Together, these rearrangements lead to enhanced engagement of the GluA2 subunit by CNIH2 and γ8.

### Counter-rotations of the auxiliary subunit pairs promote 2-fold symmetry

Expansion of the channel, triggered by M3 gate opening, transmits to both pairs of auxiliary subunits, leading to global structural changes (**Fig. 2d**): The TARP extracellular beta-sheet segment bends away from the pore axis; its TMD sector undergoes a right-handed twist around its vertical axis, while CNIH2 turns left-handedly by ~4° (**Fig. 2d-f**). As a net result, the GluA2 side of the channel (between the GluA2 M1 and M4 helices) widens on activation, while the GluA1 side (between GluA1 M1 and M4) contracts (**Extended Data Fig. 8c, d**), contributing to the two-fold symmetry-switch upon opening. The counter-rotations of the two auxiliaries are likely driven by their different topologies, and is of major consequence, as the left-handed twist of CNIH2 follows the rotation of the channel and thereby will stabilize this active conformation, with an open pore.

Moreover, rotations of the auxiliary subunits are accompanied by tilts in the vertical plane, bringing them into closer reach of cytoplasmic portions of the active receptor (**Supplementary Video 2**). We observe formation of contacts between the γ8 M4 helix and the start of the GluA1 M1/2 cytoplasmic loop (**Fig. 2e**, inset in **Extended Data Fig. 8c**), while the cytoplasmic part of the CNIH2 M2 helix tilts towards the pore (**Fig. 2f**). This behaviour is also evident by normal mode analysis (**Supplementary Video 3**) ^33,34^, where both twisting and pivoting of CNIH2 is apparent in a number of modes. Hence, activation facilitates interaction of both auxiliaries with the AMPAR intracellular portion, which is currently unresolved.

### Mechanism of CNIH2 modulation

Dictated by their architecture, CNIHs will exert AMPAR modulation exclusively at the TMD or cytoplasmic levels. Three conserved phenylalanines (Phe3, −5, −8) slot CNIH2 (and CNIH3^17^) into the upper tier of the A’C’ binding site (**Fig. 1g, i**). When mutated individually to leucine, all three mutants weakened CNIH2’s modulation of GluA2 kinetics, and reduced the equilibrium response, but did not affect complex formation or receptor trafficking (**Fig. 3a, Extended Data Fig. 9a-d**). Simultaneous mutation of all three residues to leucine reduced trafficking and, consequently, gating modulation (**Fig. 3a, Extended Data Fig. 9a-d**), highlighting this interaction as an important modulatory site.

**Fig. 3:**
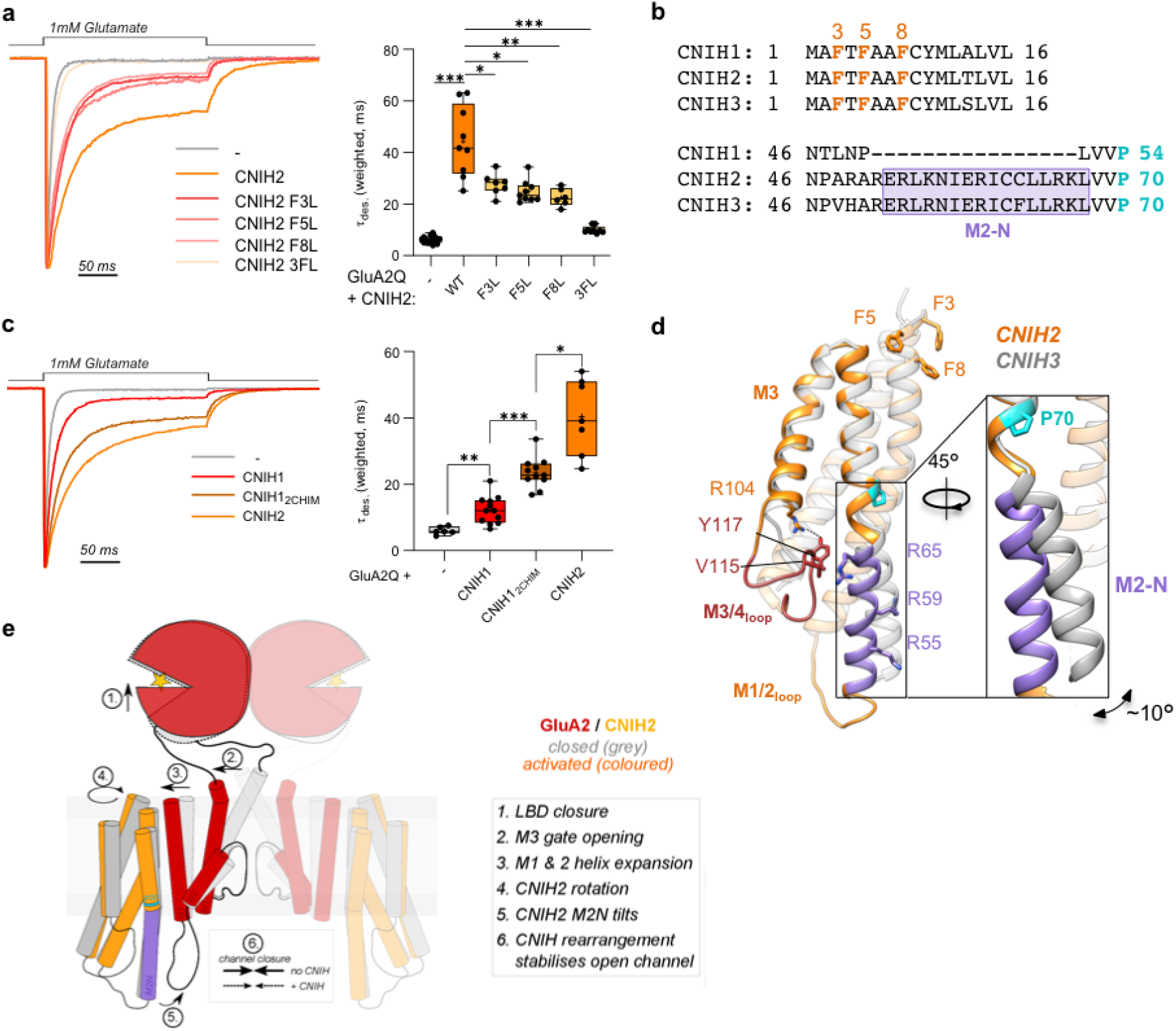
Mechanism of AMPAR modulation by CNIH2. **a**, Phe to Leu mutations of the CNIH2 N-terminus reduces modulation of GluA2Q desensitization kinetics: weighted τ_des_ (ms), mean ± SEM – GluA2 alone: 6.16 ± 0.35, n=15; CNIH2: 44.32 ± 4.58, n=9; F3L: 27.73 ± 1.58, n=7; F5L: 24.85 ± 1.43, n=9; F8L: 22.57 ± 1.40, n=6; 3FL: 10.07 ± 0.46, n=9; Welch’s ANOVA with Dunnett’s multiple comparisons test: W(5,17.83) = 85, p<0.0001. **b**, Sequence alignment of CNIH1-3 (from mouse), highlighting the conserved phenylalanines (F3, F5, F8) in the N-terminal region and the region around M2-N, where CNIH1 lacks 16 residues. Pro70 is shown in cyan. **c**, The CNIH2 M2-N helix contributes to modulation of GluA2 kinetics, demonstrated by a gain-of-function when transplanted onto the weakly modulating CNIH1: weighted τ_des_ (ms), mean ± SEM – GluA2 alone: 6.03 ± 0.47, n=6; CNIH1: 12.22 ± 1.20, n=12; CNIH1_2chimera_: 23.48 ± 1.39, n=11; CNIH2: 40.45 ± 4.32, n=7; Welch’s ANOVA with Dunnett’s multiple comparisons test: W(3,15.30) = 64.09, p<0.0001. **d**, Superposition of CNIH2 (orange) and CNIH3 (grey; PDB: 6PEQ), using the top of their M1 and M2 helices for alignment. CNIH2 M2-N (purple) kinks at Pro70 (cyan) and diverges from CNIH3 by ~ 10° (inset); N-terminal phenylalanines are also indicated. An interaction with the M3/4 loop through Val115 and Arg65 is shown, Arg55 and Arg59 project toward the pore axis. **e**, Molecular mechanism underlying CNIH2 modulation of the AMPAR.

However, the N-terminus, including Phe3, −5, −8, is conserved in CNIH1 (**Fig. 3b**), which also associates with the receptor and traffics to the cell surface, but modulates gating only weakly (**Fig. 3c, Extended Data Fig. 9a-c, e**) ^28,35^. Yet, CNIH1 lacks an elongated N-terminal segment in the M2 helix (termed ‘M2-N’) present in both modulatory subunits, CNIH2 and CNIH3 (**Fig. 3b, d**). M2-N is formed by a kink in M2, created by a conserved proline (Pro70), that ‘uncouples’ M2-N from the rest of the helix, thus enabling its tilting towards the pore on activation (**Fig. 2f, Supplementary Videos 2, 3**). Structural flexibility of M2-N is exemplified by its different kink angle between CNIH2 and CNIH3 (PDB 6PEQ). The CNIH2 M3/4 loop, with a potential cholesterol binding site in its vicinity (**Extended data Fig. 7f**), straddles M2-N, thereby impacting its dynamics (**Fig. 3d**).

To test the role of M2-N in AMPAR modulation, we inserted this segment from CNIH2 into CNIH1. The resulting CNIH1 chimera strongly gained modulatory activity, substantially slowing GluA2 desensitization and increasing the equilibrium response (**Fig. 3c, Extended data Fig.9e**), thus highlighting a critical role for CNIH2 interactions with cytoplasmic receptor elements. Given the presence of positively charged side chains projecting from M2-N towards the negatively charged M1/2 receptor loops (**Fig. 3d**), these two cytosolic elements are strong candidates to interact and enable this regulation.

## Conclusion

Based on these data, we propose the following mechanism underlying CNIH2 modulation of AMPARs (**Fig. 3e**): Association with CNIH2 stabilises the active-state conformation of pore helices, through tight local contacts between Phe3,-5,-8 and gate-surrounding receptor residues. Opening of the M3 gate triggers a left-handed rotation of CNIH2, that promotes the open state by stabilising the expanded A’C’ site helices and M2 pore loop, with a dilated pore. Full modulation requires additional contacts between M2-N and cytosolic receptor elements, which may further maintain the dilated (‘active’) channel conformation, slowing relaxation of the rotated upper regions back to the closed conformation. This scheme accounts for residual modulation by CNIH1, which supports the active state locally through its conserved N-terminus, but in lacking M2-N, is unable to maintain the fully active-state conformations strongly.

Dictated by their position in the octameric complex and by topological differences, these two auxiliary subunits act in concert to modulate AMPARs. While TARPs engage the LBDs and the M1&M3 gating linkers through their extracellular portion, CNIH2 &3 act through their distinct helical rearrangements and, now resolved, intracellular M1/2 loop element. These appendages, unique to each auxiliary subunit, stabilise transient gating conformations, and determine the rotational behavior of CNIH2 and γ8 to enact global structural changes. These mechanisms combine to build a relatively slowly deactivating AMPAR complex, providing hippocampal pyramidal neurons with the required synaptic properties to temporally coinciding inputs.

## Acknowledgements

We thank the Greger lab, Beatriz Herguedas, James Krieger, and Jan-Niklas Dohrke for comments on the manuscript; James Krieger and Jan-Niklas Dohrke for discussion, Bianka Köhegyi for help with EM imaging, and Maria Carvalho for assistance at early stages of this project. We are grateful to LMB scientific computing and the EM facility for support, Paul Emsley for help with model building, Takanori Nakane for helpful comments with Relion 3.1, and Rangana Warshamanage for helping with EMDA EM-map processing. We also acknowledge Diamond Light Source for access and support of the Cryo-EM facilities at the UK national electron bio10 imaging centre (eBIC), proposal EM17434, funded by the Wellcome Trust, MRC and BBSRC. This work was supported by grants from the Medical Research Council (MC_U105174197) and BBSRC (BB/N002113/1) to I.H.G.

## Author contributions

IHG conceptualized and supervised the study, and wrote the paper with input from JFW. DZ performed protein purification, cryo-EM data collection, data processing and model building. JFW and OC designed and performed electrophysiological experiments. PMM and JFW performed confocal imaging and data analysis.

## Data and materials availability

Cryo-EM density and PDB models have been submitted to the PDB and EMDB.

## Methods

### cDNA constructs

All cDNA constructs were produced using IVA cloning ^37^. To achieve heteromeric AMPAR expression, GluA1 (rat cDNA sequence, flip isoform) was co-expressed with GluA2 (rat cDNA sequence, flip isoform, R/G edited, Q/R edited), expressed from the pRK5 vector. For TARP-containing complex recordings, GluA2 was expressed in a tandem configuration (denoted GluA2_γ8) with TARP γ8 (rat cDNA sequence) by cloning the TARP γ8 coding sequence (Glu2 - Lys419) at the C-terminus of the GluA2 coding sequence (Val1 to Ser839), in the pRK5 vector, separated by a Gly-Gly-Ser-Gly-Ser-Gly linker sequence. pN1-EGFP (Clontech) was used for visualisation of transfected cells. CNIH2 (rat) was expressed either from a stable cell line (heteromeric AMPAR recordings, see below), or from either a pRK5 or a pBOS vector, with an HA-tag at the extreme C-terminus. A chimera of CNIH1 and CNIH2 (CNIH1_2CHIM_) was made by inserting 16 amino acids from CNIH2 (51-RERLKNIERICCLLRK-66) into CNIH1 at position P50-L51.

### Organotypic slice preparation

All procedures were carried out under PPL 70/8135 in accordance with UK Home Office regulations. Experiments conducted in the UK are licensed under the UK Animals (Scientific Procedures) Act of 1986 following local ethical approval.

Organotypic slice cultures were prepared as described previously ^38^. Briefly, hippocampi from P6-8 C57/Bl6 mice were isolated in high-sucrose Gey’s balanced salt solution containing (in mM): 175 sucrose, 50 NaCl, 2.5 KCl, 0.85 NaH_2_PO_4_, 0.66 KH_2_PO_4_, 2.7 NaHCO_3_, 0.28 MgSO_4_, 2 MgCl_2_, 0.5 CaCl_2_ and 25 glucose at pH 7.3. Hippocampi were cut into 300 μm thick slices using a McIlwain tissue chopper and cultured on Millicell cell culture inserts (Millipore Ltd) in equilibrated slice culture medium (37°C/5% CO_2_). Culture medium contained 78.5% Minimum Essential Medium (MEM), 15% heat-inactivated horse serum, 2% B27 supplement, 2.5% 1 M HEPES, 1.5% 0.2 M GlutaMax supplement, 0.5% 0.05 M ascorbic acid, with additional 1 mM CaCl_2_ and 1 mM MgSO_4_ (all from Thermo Fisher Scientific; Waltham, MA). Medium was refreshed every 3–4 days. Recordings were performed at 7-14 days *in vitro*.

### Recombinant cell electrophysiology

For heteromeric AMPAR experiments, suspension HEK-Expi293F™ cells (ThermoFisher Scientific) were cultured in Expi293™ expression media (Gibco) and were transfected using PEI Max with a DNA to PEI ratio at 1:3. GluA1, GluA2 (with or without TARP γ8 tethered via a flexible linker, as described before ^15^ and EGFP plasmids were transfected at a 2:8:1 stoichiometry to aid heteromeric receptor production, and identification of transfected cells. For homomeric AMPAR experiments, GluA1 or GluA2Q (pIRES2-mCherry or pIRES2-EGFP) and CNIH2 (pRK5 or pBOS) plasmids were transfected at a 1:2 ratio using Effectene (QIAGEN) into adherent HEK293T cells (*ATCC: Cat# CRL-11268, RRID: CVCL_1926, Lot 58483269: identity authenticated by STR analysis, mycoplasma negative)*. Cells were cultured at 37°C and 5% CO_2_ in DMEM (Gibco; high glucose, GlutaMAX, pyruvate, Cat#10569010) supplemented with 10% foetal bovine serum (Gibco) and penicillin/streptomycin. Transfected cells were recorded from 36 hours post-transfection. Cells were plated on poly-L-lysine coated glass coverslips approximately 12 hours before recording, to allow outside-out patch recording and 30 μM 2,3-dioxo-6-nitro-1,2,3,4-tetrahydrobenzo[f]quinoxaline-7-sulfonamide (NBQX; Tocris, Cat#1044) was added to media post-transfection to avoid AMPAR-mediated toxicity.

Borosilicate glass electrodes (1.5mm o.d., 0.86mm i.d., Science Products GmbH), pulled with a PC-10 vertical puller (Narishige) with tip resistance of 2 to 6 MΩ, were filled with internal solution containing (in mM) CsF (120), CsCl (10), EGTA (10), HEPES (10), and spermine (0.1), adjusted to pH 7.3 with CsOH. Surface membrane patches were acquired by gentle excision after achieving the whole-cell patch clamp configuration. Patches were voltage clamped at −60 mV, and subject to fast application of 1 mM L-glutamate using two-barrel theta glass tube controlled by a piezoelectric translator (Physik Instrumente or Burleigh), allowing solution exchange in around 200 μs (open tip response). Signals were acquired using the MultiClamp 700B or Axopatch 200B amplifier (Axon Instruments), digitised using a Digidata 1440A interface and recorded with pClamp10 (Molecular Devices). Extracellular solution contained (in mM) NaCl (145), KCl (3), CaCl_2_ (2), MgCl_2_ (1), glucose (10), and HEPES (10), adjusted to pH 7.4 using NaOH. 20 μM IEM 1925 dihydrobromide (Tocris, Cat#4198) was added to the extracellular solution to limit the contribution of homomeric AMPAR complexes to heteromeric recordings.

Current/voltage relationships were recorded for AMPAR responses with holding potentials of −100 mV to +100 mV in 20 mV steps. To ensure heteromeric receptor analysis, patches with a rectification index of less than 0.5 were discarded. Recordings were not corrected for the liquid junction potential. Rectification index was calculated as follows:

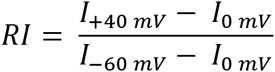

Desensitization entry was determined from the first 150-200 ms after the peak response, which was fitted with a two-exponential function to obtain the (weighted) time constant. ‘Equilibrium’ responses were denoted as the percentage of peak current remaining after 200 ms.

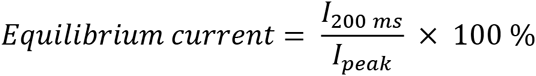

Signal acquisition and data analysis were performed using pClamp10. Neuronal cell electrophysiology

Cultured organotypic hippocampal slices were submerged in artificial cerebrospinal fluid (aCSF) containing (in mM): 125 NaCl, 2.5 KCl, 1.25 NaH_2_PO_4_, 25 NaHCO_3_, 10 glucose, 1 sodium pyruvate, 4 CaCl_2_ and 4 MgCl_2_ at pH 7.3 and saturated with 95% O_2_/5% CO_2_. Neuronal surface patches were acquired from the cell soma as detailed above for recombinant cell patches, using borosilicate pipettes containing (in mM): 135 CH_3_SO_3_H, 135 CsOH, 4 NaCl, 2 MgCl_2_, 10 HEPES, 4 Na_2_-ATP, 0.4 Na-GTP, 0.15 spermine, 0.6 EGTA, 0.1 CaCl_2_, at pH 7.25. Agonist was applied to neuronal patches using a piezo-driven theta-barrelled perfusion setup, with aCSF containing (in mM): 140 NaCl, 3.5 KCl, 1 MgCl_2_, 2.5 CaCl_2_, 10 HEPES, 10 glucose, 1 sodium pyruvate, 2 NaHCO_3_, at pH 7.3. 100 μM D-APV (Tocris, Cat #0106) was added to the extracellular solution of both barrels to prevent the contribution of NMDA receptor currents. 1 mM L-glutamate was added to one barrel for fast-agonist application (as above). A subset of hippocampal CA1 pyramidal neuron patches were recorded using the same extra- and intracellular solutions as recombinant cell electrophysiology, and no difference in kinetic parameters analysed was observed between recording configurations, therefore these datasets were pooled and presented together.

CA1 and CA3 pyramidal, and dentate gyrus (DG) granule cells were identified by cell morphology and location in tissue. *Stratum radiatum* interneurons (IN) were selected for analysis as a population of interneuronal cells that can be identified by location, rather than requiring a genetic driver line, and consisted of neurons located between *stratum pyramidale* and *stratum lacunosum-moleculare* in the CA1 region of organotypic slices. Cell identity was not confirmed by morphological or other analysis *post-hoc*, and therefore likely consists of a heterogeneous combination of IN cell types.

### Generation of CNIH2-1D4-HA stable cell line

A CNIH2-1D4-HA stable expression cell line was generated using a lentiviral expression system following an established protocol ^39^. A 1D4 tag followed by an HA tag were included at the extreme C-terminus of rat CNIH2 with a TEV cleavage site, separated by a ‘GGS’ linker sequence. The CNIH2 gene together with the two tags were synthesized and cloned into pHR vector. pHR-CNIH2-ID4-HA was co-transfected with psPAX2 and pMD2.G into HEK293T Lenti-X cells (Takara/Clontech, Cat# 632180). 72 hours after transfection, lentiviral particles were harvested from the media and used to infect HEK-Expi293F cells. 48 hours after infection, fresh Expi293™ expression media (Gibco) was added, cells were surface-labelled with an APC-conjugated anti-HA antibody (Miltenyi Biotec, Cat# 130-098-404, 1:50) and sorted by flow cytometry, collecting HA-positive and DAPI-negative cells. Positive cells were then scaled up for A1/A2_γ8/C2 complex expression.

### Expression and purification of A1/A2_γ8/C2

Constructs of FLAG-tagged GluA1(flip) and GluA2(flip)-TARPγ8-eGFP tandem (pRK5) used in this paper are the same as previously reported ^15^. GluA1 and GluA2-TARPγ8 were co-transfected into the CNIH2-1D4-HA stable expression cell line at a ratio of 1:1. To prevent AMPA-mediated excitotoxicity, AMPAR antagonists ZK200775 (2 nM, Tocris, Cat# 2345) and kynurenic acid (0.1 mM, Sigma, Cat# K335-5G) were added to the culture medium. 36-44 hours post-transfection, cells were harvested and lysed for 3 hours in lysis buffer containing: 25 mM Tris pH 8, 150 mM NaCl, 0.6 % digitonin (w/v) (Sigma, Cat# 300410-5G), 5 μM NBQX, 1 mM PMSF, 1× Protease Inhibitor (Roche, Cat# 05056489001). Insoluble material was then removed by ultracentrifugation (41,000 rpm, 1 hour, rotor 45-50 Ti) and the clarified lysate incubated with anti-GFP beads for 3 hours. After washing with glyco-diosgenin (GDN) (Anatrace, Cat# GDN101) buffer (25 mM Tris pH 8, 150 mM NaCl, 0.02% GDN) the protein was eluted from the beads by digestion with 1 mg/ml 3C protease at 4 °C overnight. Eluted fractions were incubated with FLAG beads (Sigma, Cat# A2220) for 1.5 hours and washed 3 times with GDN buffer. Finally, the complex was eluted using 0.15 mg/ml 3×FLAG peptide (Millipore Cat# F4799) in GDN buffer. Eluted fractions were pooled and concentrated to ~3 mg/ml for cryo-EM grid preparation.

### Cryo-EM grid preparation and data collection

Cryo-EM grids were prepared using a FEI Vitrobot Mark IV. For the resting state complex, protein was incubated with 100 μM NBQX for at least 30 min on ice before freezing. For the active state complex, protein was first incubated with 300 μM Cyclothiazide (CTZ, Tocris, Cat# 0713) for at least 30 min on ice and then quickly mixed with 1 M L-glutamate stock solution to a final concentration of 100 mM prior to loading onto the grids. Quantifoil Au 1.2/1.3 grids (300 mesh) were glow-discharged for 30 s at 0.35 mA before use. 3 μl sample was applied to the grids, blotted for 4.5-5 s at 4 °C with 100 % humidity and plunge-frozen in liquid ethane. Cryo-EM data were collected on an FEI Titan Krios operated at 300 kV, equipped with a K3 detector (Gatan) and a GIF Quantum energy filter (slit width 20 eV). Movies at 1.5-2.5 μm underfocus were taken in counting mode with a pixel size of 1.1 Å/pixel. A combined total dose of 50 e/Å^2^ was applied with each exposure and 50 frames were recorded for each movie. Two datasets of resting state were collected using SerialEM with 10462 movies in total, and two datasets of active state were collected using EPU2 with 8246 movies in total.

### Cryo-EM data processing and model building

Dose-fractionated image stacks were first motion-corrected using MotionCor2 ^40^. Corrected sums were used for CTF estimation by GCTF ^41^. All further data processing was performed with RELION 3.1 ^42^. Automatic particle picking was performed using a Gaussian blob and particles were binned to 4.4 Å/pixel and extracted in a box of 80 pixels. 2 to 3 rounds of 2D classification were carried out to remove particles not showing AMPAR-like features. For the following 3D classification, emd-20332 was used as initial model to further eliminate low-quality particles. Following data clean-up, particles were re-centered, scaled up to 2.2 Å/pixel and re-extracted in a box of 160 pixels. Another 3D classification focused only on the LBD-TMD and C2 symmetry was applied on the re-extracted data set with EMD-20330 as an initial model. After this round of classification, CNIH2 containing and CNIH2-free particles were separated into two datasets with 3D refinement applied on each set. Refined particles were scaled to the original 1.1 Å/pixel size, and refined with C2-symmetry followed by post-processing. Four datasets were processed independently in the beginning, and particles at the same states were joined together before CTF refinement and Bayesian polishing. The final refinement was carried with side-splitter to achieve maps at 3.2 Å (CNIH2 containing) and 3.6 Å (CNIH2 free) resolution in the resting state and 3.7 Å (CNIH2 containing) in the active state (individually based on the standard Fourier shell correlation (FSC) 0.143). To further improve map resolution, we applied masked refinement on TMD or LBD region independently by continuing old run from the last iteration of the global LBD-TMD refinement. Local resolution was also estimated by RELION3.1. To help with model building, we used EMDA (https://www2.mrc-lmb.cam.ac.uk/groups/murshudov/content/emda/emda.html) to generate composite maps of LBD-TMD from the mask refinement. However, the density observed at the CNIH region was weaker than that of the receptor. In order to further enhance the density for model building, symmetry expansion was applied on the aligned particles from TMD reconstruction. A masked classification without alignment was then performed to select CNIH stable particles. To process the active state data, a smaller mask and higher T value were required for the focused classification. The following focused refinement was found to improve the density of the CNIH region in both resting and active states. For reconstruction of the NTD in the resting state, after scaling particles back up to 2.2 Å/pixel, 3D classifications against the full-length receptor were performed and particles with a stable NTD signal were selected and subjected to masked refinement with a soft mask placed only on the NTD. CTF refinement and Bayesian polishing were also preformed to attain a final 3.5 Å map.

Model building and refinement were performed using Coot ^43^, REFMAC5 ^44^ and PHENIX ^45^ real-space refinement. For modelling, the GluA1/A2-γ8 complex (PDB 6QKC and 6QKZ) and CNIH3 in GluA2-CNIH3 complex (PDB 6PEQ) were used as starting points. The individual coordinates were first rigid-body fitted into the map using UCSF chimera (http://www.rbvi.ucsf.edu/chimera) and then automatically refined by REFMAC5. After this, manual refinement through Coot was performed, followed by PHENIX real-space refinement, to further refine the geometry. External restraints generated from LBD crystal structure 3TKD were used to further improve the model quality in the LBD region. For lipid building, EMDA bestmap was used to enhance the lipid signal in the maps. Model validation was performed with MolProbity ^46^. All graphics figures in the paper were prepared using UCSF Chimera or PyMOL (http://www.pymol.org). Pore radius was calculated using a plugin version of HOLE ^47^ in Coot.

### Normal mode analysis

Normal modes were calculated and visualized using the ANM server ^34^ and the *ProDy* API ^48^ In both cases, the protein was treated as an elastic network model as previously described ^33^. Briefly, each residue is represented as one bead placed at the Cα position and interactions were treated as uniform harmonic springs within a cutoff distance of 15 Å. This simple representation allows fast analytic calculation of the most energetically favourable modes of motion via normal mode analysis. Calculations were facilitated using the *SignDy* module within *ProDy* ^49^, which enabled easier alignment of the active and resting structures and comparisons of their modes to each other and the transition between them, which were calculated using the correlation cosine overlap between vectors. Movies were created using PyMOL version 1.8.2.0 and Fiji ^50^.

### Flag Immunoprecipitation of CNIH homologues and CNIH2 mutants

All CNIH homologues, the CNIH12 chimera and CNIH2 mutants were HA-tagged at the extreme C-terminus and cloned into the pRK5 vector. FLAG-tagged GluA2 was co-transfected alongside CNIH constructs. 10 ml of HEK-Expi293F cells was used for each transfection. 40 hours post-transfection, cells were harvested and resuspended in 1 ml lysis buffer (25 mM Tris pH 8, 150 mM NaCl, 0.6 % digitonin (w/v)). After a 2 hour incubation, cell lysate was centrifuged at 21000 × g for 30 min and the clarified supernatant was taken as the input sample. 10 μl FLAG beads was added to the input and incubated for 1.5 hours. Following incubation, FLAG beads were pelleted and washed 5× with GDN buffer (25 mM Tris pH 8, 150 mM NaCl, 0.02% GDN). To elute samples the beads were then boiled in SDS sample buffer for 5 mins and run on a 4-12% Bis-Tris gel (Invitrogen, Cat# NW04127BOX). Rabbit GluR2 polyclonal antibody (Millipore, Cat# AB1768-l) and rabbit HA antibody (Sigma, Cat #6908) were used to detect the GluA2 and CNIHs signal, respectively.

### Immunostaining

Suspension HEK-Expi293F™ cells were transfected with GluA2Q (pRK5) and CNIH (pRK5) at a stoichiometry of 1:1 and settled on poly-L-lysine coated 12 mm glass coverslips (Corning) 36 hours later. Live-labelling of surface CNIH was performed at 48 hours post-transfection by incubation with rabbit anti-HA (Sigma, Cat# H6908; 1:200) for 20 min at room temperature (RT) in Expi293™ expression media (Gibco). Cells were washed 3× in medium and once in PBS before fixation in 4% paraformaldehyde and 4% sucrose for 10 min at RT. Subsequently, cells were washed in PBS before permeabilisation in 0.1% Triton X-100 (Fisher Bioreagents) and blocking in 1% bovine serum albumin (BSA; Fisher Bioreagents) and 10% normal goat serum (NGS; Sigma) in PBS for 30 min. Permeabilised cells were then incubated sequentially in primary and secondary antibodies prepared in 1% BSA and 10% NGS for 2 hours at RT and washed in PBS after each incubation. Total CNIH was detected using mouse anti-HA (BioLegend, Cat# 901501; 1:500) and total GluA2 was detected using guinea pig anti-GluA2 (Synaptic Systems, Cat# 182 105; 1:500), followed by incubation in corresponding secondary antibodies: goat anti-rabbit IgG Alexa Fluor (AF) 568 (Invitrogen, Cat# A-11036; 1:500), goat anti-mouse IgG AF 488 (Invitrogen, Cat# A-11029; 1:500) and goat anti-guinea pig IgG AF 647 (Invitrogen, Cat# A-21450; 1:500). Coverslips were mounted in ProLong Diamond Antifade Mountant (Invitrogen) and left to cure in the dark for 48 hours at RT.

Images were acquired with a Leica TCS SP8 confocal inverted microscope using Leica Application Suite X software. Z-stacks (5x 1 μm steps) of whole cells were acquired using a 63x oil-immersion objective. 6 images were taken per coverslip and repeated from 3 independent preparations. Equivalent laser power settings were applied across all experimental conditions. Z-stacked images were averaged and the fluorescence intensity was measured from individual cells using ImageJ (Fiji) and normalised for cell area and background fluorescence. Brightness and contrast settings were kept consistent across all images.

