## Supplementary Information for "Gating transitions and modulation of a hetero-octameric AMPA glutamate receptor"

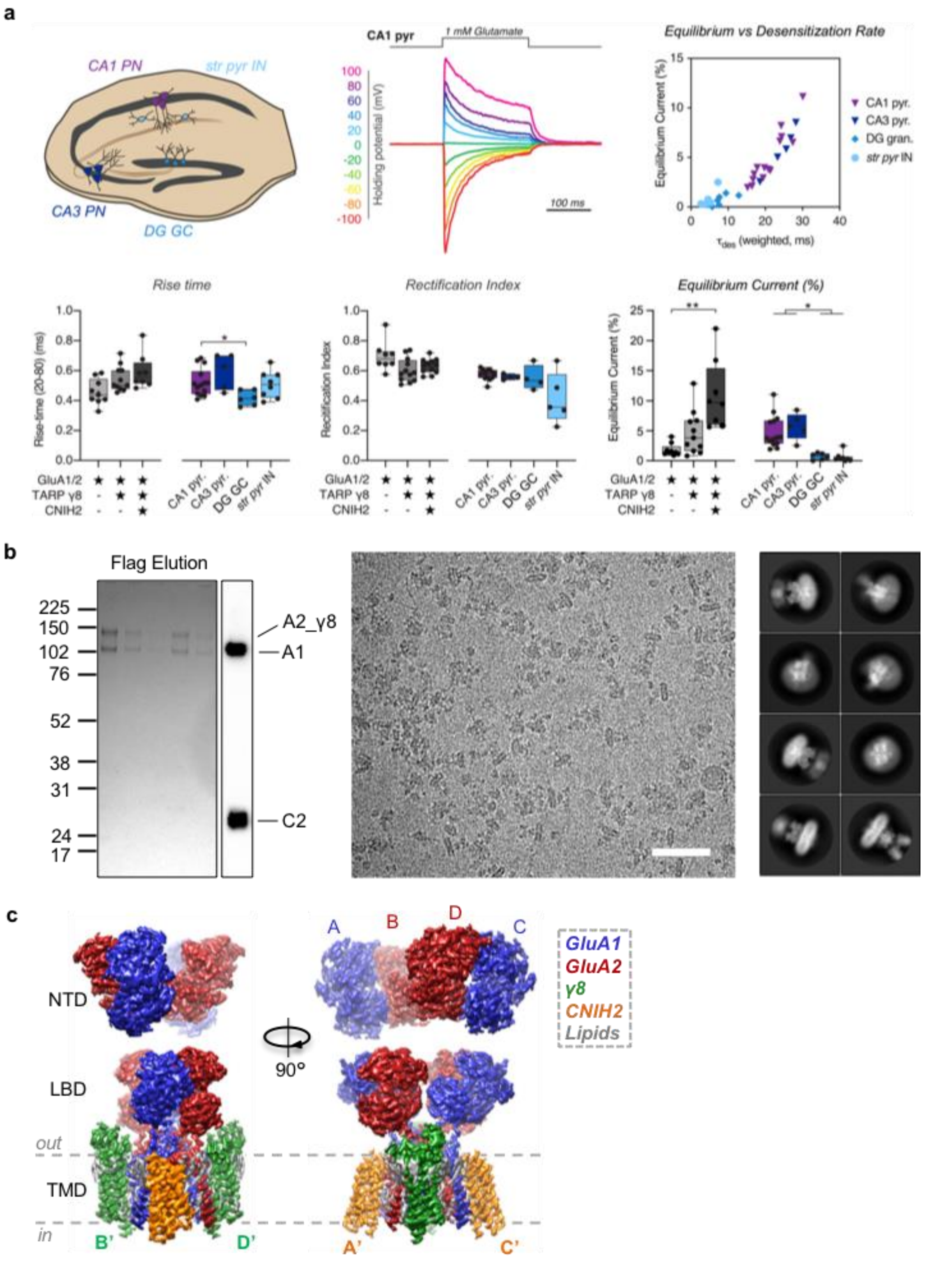


**Extended Data Figure 1 | Properties of neuronal and recombinant AMPAR complexes. a**, Electrophysiological properties of neuronal and recombinant AMPAR complexes. **Top left**: Hippocampus schematic indicating selected cell types. **Bottom left**: Rise-time of fast-application glutamate responses of recombinant and neuronal AMPAR patches (20-80 % rise time (ms) - *Recombinant receptors:* GluA1/2: 0.46 ± 0.03, n=9; +γ8: 0.55 ± 0.03, n=11; +γ8+CNIH2: 0.59 ± 0.04, n=8. *Neuronal receptors:* CA1 pyramidal: 0.52 ± 0.02, n=14; CA3 pyramidal 0.60 ± 0.05, n=5; DG granule cell: 0.42 ± 0.02, n=6; CA1 stratum pyramidale interneurons: 0.51 ± 0.03, n=8; Welch's ANOVA with Dunnett's multiple comparison tests – Recombinant: W(2,15.11) = 4.25, p=0.03; Neurons: W(3,12.12) = 5.40, p=0.014). **Top middle**: Example trace of rectification index (RI) recording from CA1 pyramidal neuron normalized to -100 mV peak amplitude. **Bottom middle**: Quantified RI from recorded surface patches (*Recombinant receptors:* GluA1/2: 0.70 ± 0.04, n=8; +γ8: 0.60 ± 0.02, n=12; +γ8+CNIH2: 0.63 ± 0.01, n=12. *Neuronal receptors:* CA1 pyramidal: 0.58 ± 0.01, n=13; CA3 pyramidal 0.56 ± 0.01, n=4; DG granule cell: 0.55 ± 0.04, n=4; CA1 stratum pyramidale interneurons: 0.42 ± 0.08, n=5; Welch's ANOVA with Dunnett's multiple comparisons test – Recombinant: W(2,15.01) = 2.47, p=0.12, Neurons: W(3,7.57) = 1.5, p=0.29). **Top right**: Strong correlation between equilibrium current and desensitization rate are observed (individual neuronal patches plotted). **Bottom right**: Equilibrium current for patch responses show auxiliary protein dependent modulation and neuronal heterogeneity (% peak current - *Recombinant receptors:* GluA1/2: 1.81 ± 0.34, n=9; +γ8: 4.72 ± 1.09, n=11; +γ8+CNIH2: 10.97 ± 2.03, n=8. *Neuronal receptors:* CA1 pyramidal: 4.86 ± 0.71, n=14; CA3 pyramidal 5.78 ± 1.00, n=5; DG granule cell: 0.75 ± 0.22, n=6; CA1 stratum pyramidale interneurons: 0.59 ± 0.29, n=8; Welch's ANOVA tests with Dunnett's multiple comparisons test – Recombinant: W(2,12.09) = 11.93, p=0.002; Neurons: W(3,12.42) = 16.74, p=0.0001). **b**, Purification and cryo-EM images of the GluA1/2_γ8/CNIH2 complex. **Left**: 4-12% Bis-Tris gel stained with coomassie blue, indicating elution of A1/2_γ8/C2 complex from FLAG beads. CNIH2 expression is detected by probing for the C-terminal HA tag on western blot . **Middle**: Motion-corrected micrograph of the resting state A1/2_γ8/C2 complex (scale bar, 50 nm). **Right**: Representative 2D class averages of the resting state A1/2_γ8/C2 complex. **c**, Cryo-EM maps of the full-length AMPAR octamer, depicting the three domain layers, NTD, LBD and TMD, composed of the GluA1 (blue), GluA2 (red) heteromer associated with γ8 (green) and CNIH2 (orange)..

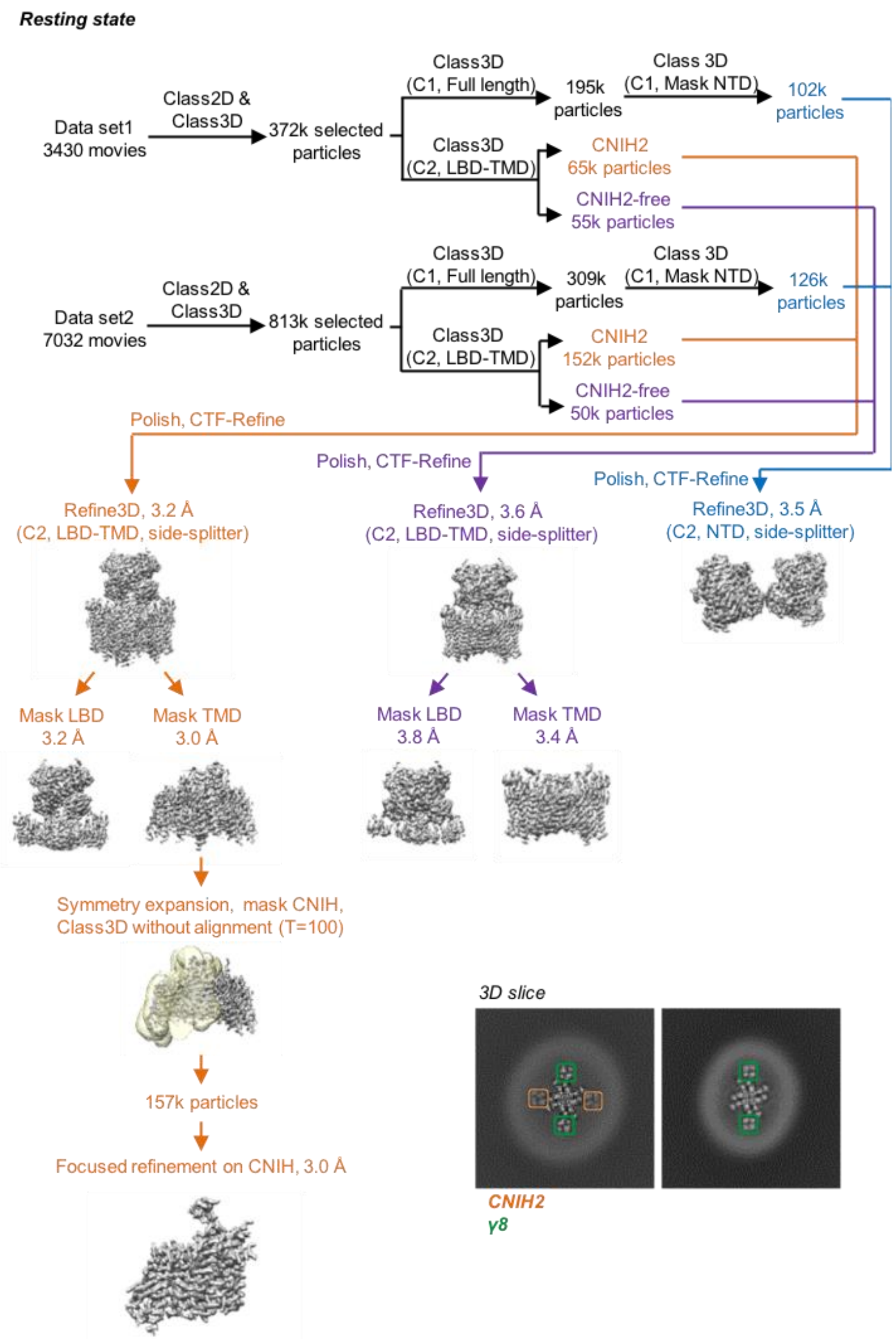


**Extended Data Figure 2 | Cryo-EM data processing workflow of the resting state A1/2_γ8/C2 complex.** Two datasets were first processed individually to remove particles lacking AMPAR features. Next, classifications focused on the LBD-TMD region were performed to separate CNIH-containing and CNIH-free particles, meanwhile classifications for full-length receptors were conducted to elucidate particles with a stable NTD signal. Subsequently, particles from the two datasets were combined together for refinement. Focused refinements were performed separately on the LBD-TMD gating core and the NTD region. To further improve the resolution, LBD and TMD are refined separately A structure of A1/2_γ8 (lacking CNIH2) was also resolved from the same dataset (containing only γ8 observed in 3D slice). CNIH density was further enhanced by applying first symmetry expansion on aligned particles from the TMD reconstruction, following by focused classification and refinement on only CNIH2 and the surrounding receptor transmembrane helices. **Inset**: Top view slices of the A1/2_γ8/C2 (left) and A1/2_γ8 (right). 3D maps at the TMD region show signal for transmembrane helices of γ8 (green) and CNIH2 (orange).

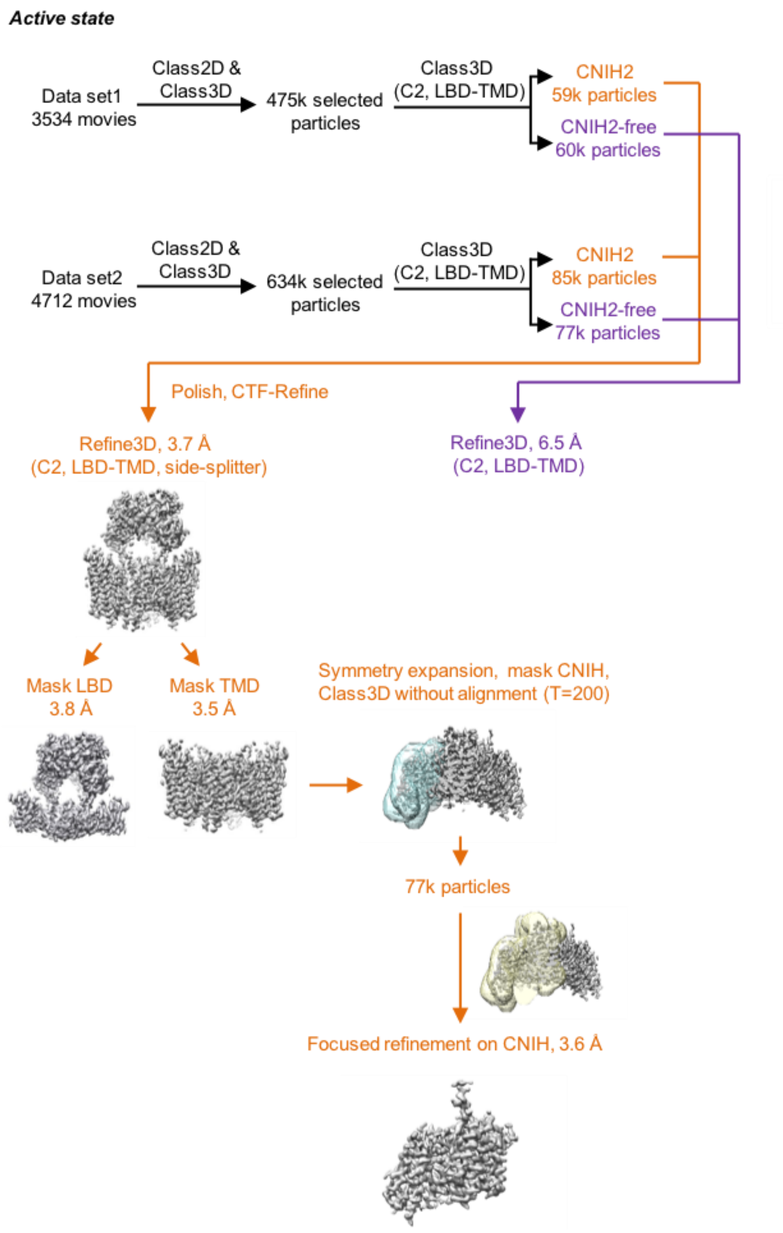


**Extended Data Figure 3 | Cryo-EM data processing workflow of the active state A1/2_γ8/C2 complex.** The overall data processing procedure for the active state complex is similar to that of resting state complexes. Focused refinement was performed on the LBD-TMD gating core and individual LBD and TMD domain layers of the receptor. CNIH density was further improved by applying first symmetry expansion on aligned particles from the TMD reconstruction, followed by focused classification on CNIH alone, and finally focused refinement on CNIH together with surrounding receptor transmembrane helices. Particles lacking CNIH2 found in these datasets were not of high enough quality to provide a high-resolution structure.

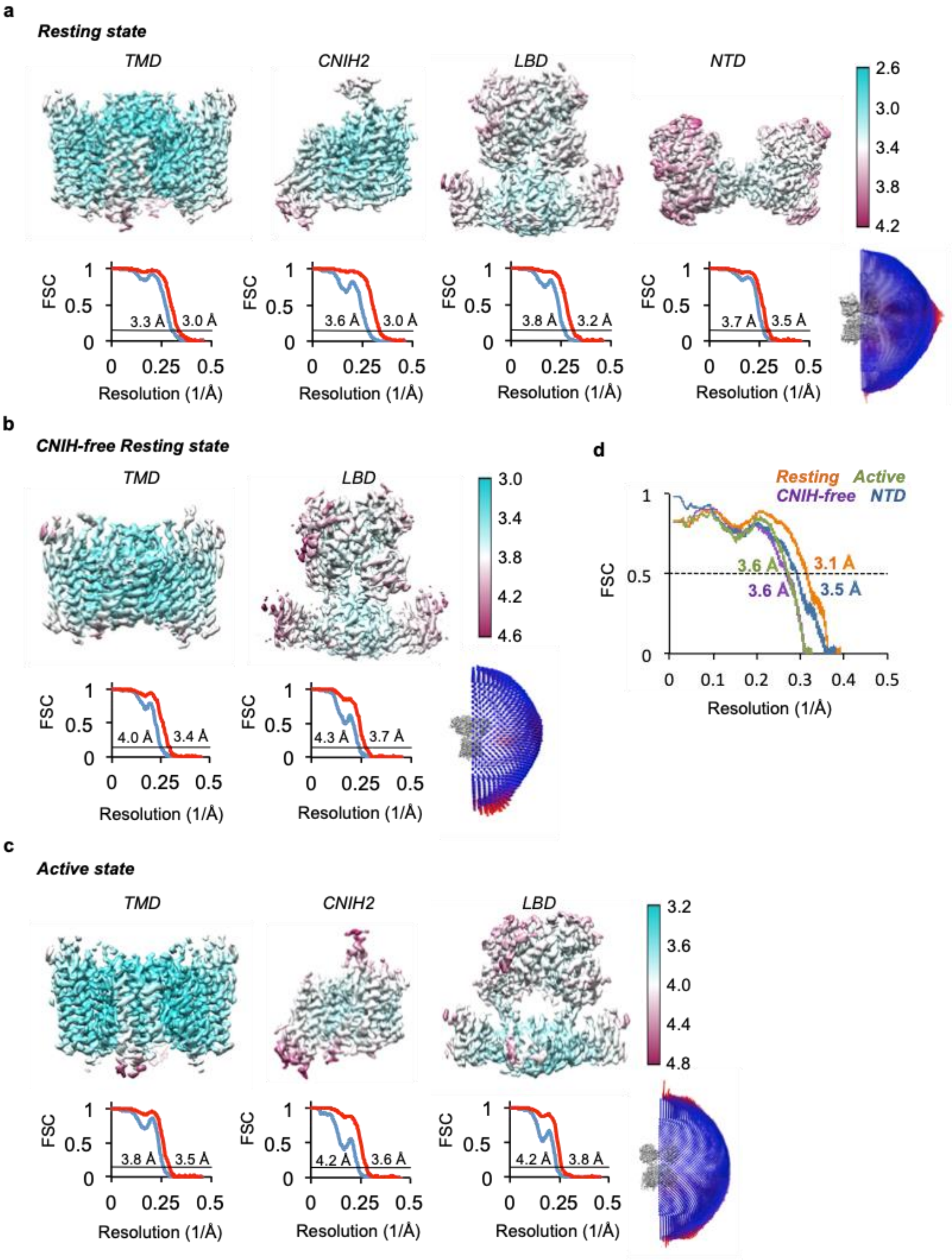


**Extended Data Figure 4 | Cryo-EM analysis of A1/2_γ8/C2 and A1/2_γ8 complexes. a**, Local resolution and Fourier shell correlation (FSC) of focused refinement maps at the TMD, CNIH2, LBD and NTD. Euler angle distribution of particles for cryo-EM reconstruction of the resting state A1/2_γ8/C2 complex. 3D maps are coloured based on local resolution estimation. Masked (red) or unmasked (blue) FSC of corresponding maps are both shown where FSC=0.143 (black line). **b**, Local resolution and FSC of focused refinements at the TMD and LBD. Euler angle distribution of particles for cryo-EM reconstruction of the resting state A1/2_γ8 complex. **c**, Local resolution, FSC of focused refinements at the TMD, CNIH2 and LBD. Euler angle distribution of particles for cryo-EM reconstruction of the active state A1/2_γ8/C2 complex. **d**, Model to map FSCs of A1/2_γ8/C2 LBD-TMD models in resting and active states, resting state NTD model and resting state A1/2_γ8 LBD-TMD model.

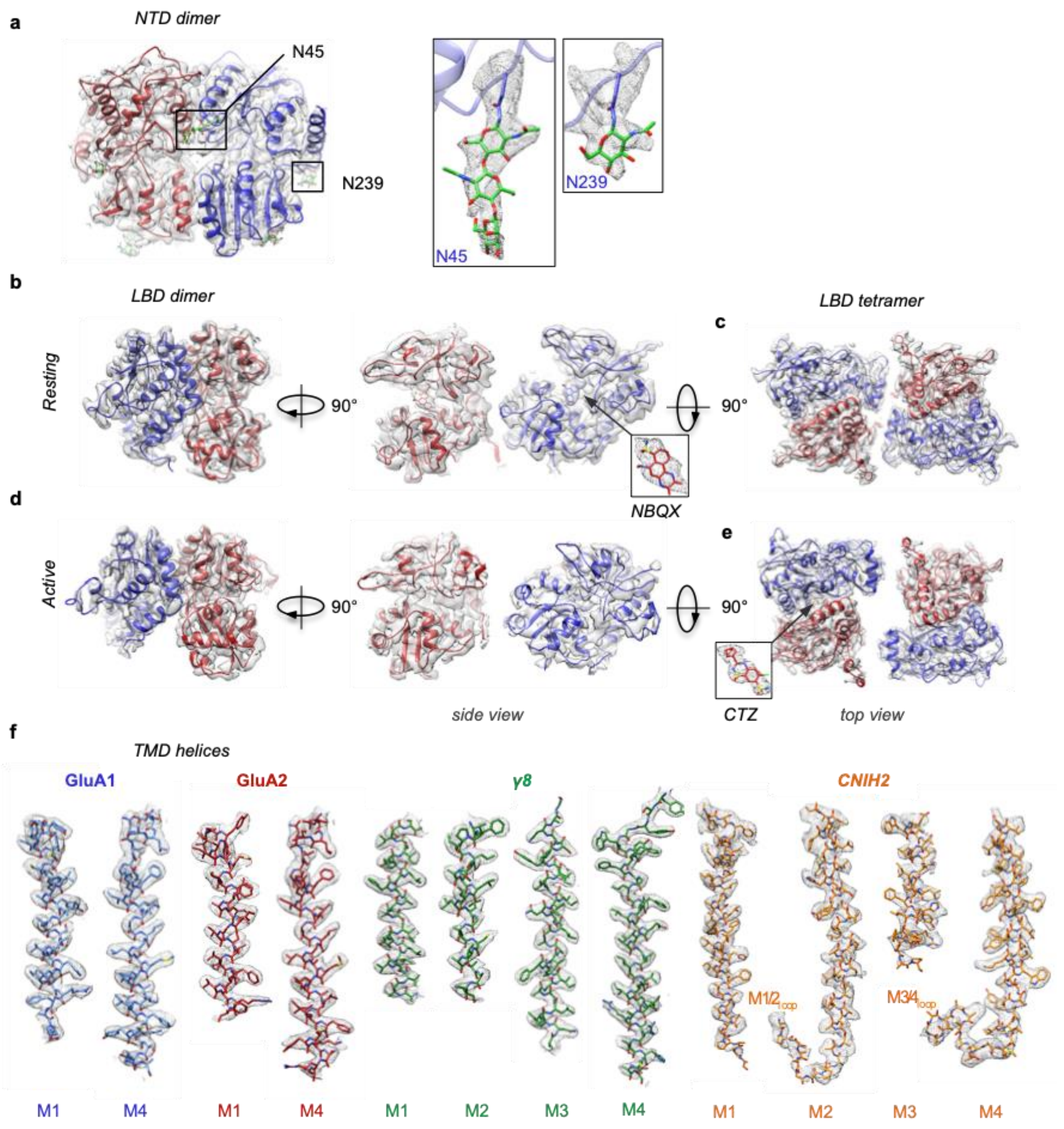


**Extended Data Figure 5 | Features of A1/2_γ8/C2 NTD and LBD layers and quality of density in the TMD region.** **a**, Cryo-EM density and model of the resting state GluA1 (blue) and GluA2 (red) NTD dimer. GluA1-specific N-linked glycans are observed at N45 and N239 (green sticks). **b**, Cryo-EM density and model of A1/2 LBD dimer in the resting state. Density and model for the competitive antagonist NBQX bound to its orthosteric site in the LBD cleft. **c**, Top view of cryo-EM density and model of A1/2 LBD tetramer in the resting state. **d**, Cryo-EM density and model of A1/2 LBD dimer in the active state demonstrating a closure of the LBD 'clamshell'. **e**, Top view of cryo-EM density and model of A1/2 LBD tetramer in the active state. Density and model of desensitization blocker cyclothiazide (CTZ) bound at the LBD dimer interface are shown in the insert. **f**, Cryo-EM density and model of transmembrane helices of A1/2_γ8/C2 in the resting state.

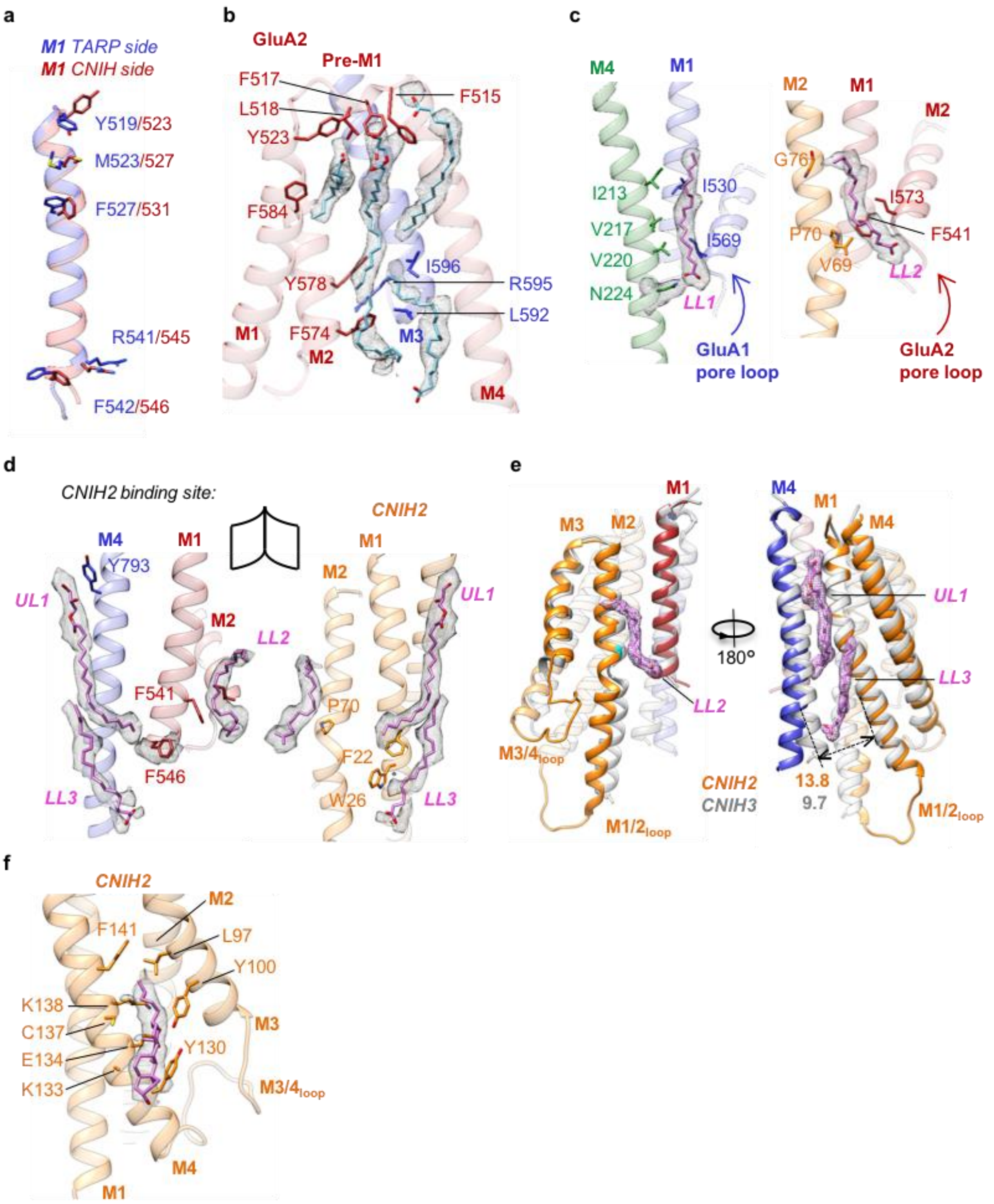


**Extended Data Figure 6 | γ8 and CNIH2 receptor binding sites and their relevant bound-lipids.** **a**, Overlay of the A'C' and B'D' binding sites, showing reorientations of five residues along the M1 helices (GluA1 M1, red; GluA2 M1, blue). These changes are likely mediated by γ8 engaging GluA1 M1. **b**, Strong lipid densities (light blue, density shown in grey mesh) line the cavity between GluA2 M1 and M4. F515, F517 and L518 from GluA2 pre-M1 interact with the lipids from the upper leaflet. Other residues from GluA2 M1, M2 and GluA1 M3 involved in these interactions are shown as stick. **c**, LL1 binds to the γ8 N224 side chain, connecting TARP-γ8 to the GluA1 M2 pore helix. LL2 bridges between CNIH2 and the GluA2 M1 and M2 helices. **d**, 'Open book' view of the A'C' binding site, displaying how the UL1, LL2 and LL3 lipids engage the receptor (left) and the CNIH2 M1 and M2 helices (right). Side chains in close proximity to lipids are shown. **e**, Superposition of CNIHs and their binding peripheral helices from the resting state A1/2_γ8/C2 (orange) and A2_C3 (grey, PDB 6PEQ) complexes. While the upper part of CNIH's M1 and M2 helices are aligned together, the lower part of CNIH2 is kinked away from the receptor relative to CNIH3, this permits the accommodation of three CNIH2 binding-relevant lipids. Distance between W26 (C2) and C811 (A1) in A1/2_γ8/C2 and W26 (C3) and C815 (A2) in A2_C3 are measured. M1 and M4 from A1/A2_γ8/C2 resting state are coloured as in Figure 1. M1 and M4 for A2_C3 (PDB 6PEQ) are coloured in grey. Three CNIH2 binding-relevant lipids LL2, LL3 and UL1 are shown as pink stick. **f**, A density modeled as cholesterol occupies the pocket between CNIH2 M3 and M4, observed after focused refinement.

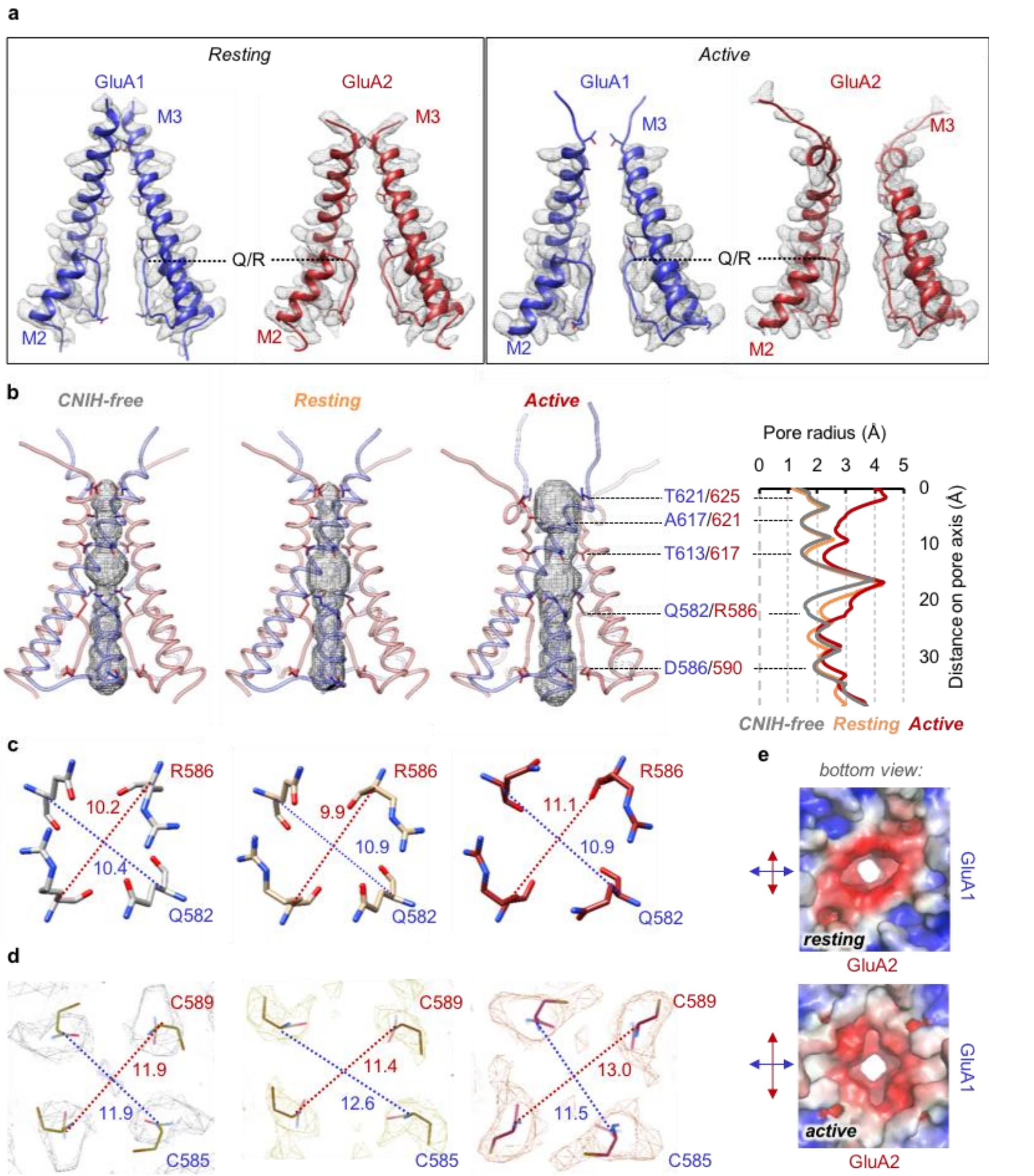


**Extended Data Figure 7 | Features of the A1/2_γ8/C2 and A1/2_γ8 conduction pore.** **a**, Density of M2/M3 gating regions and their fit against models in the resting and active state. **b**, Pore dimensions of resting state A1/2_γ8 (left) and the resting (middle) and active (right) state of A1/2_γ8/C2 depicted by space-filling representation (HOLE program) with relevant side chains indicated as sticks. A comparison of pore radius across these three structures indicates a similar diameter of the receptor gate in resting state A1/2_γ8 (grey) and A1/2_γ8/C2 (orange), with a clear expansion observed in the active state A1/2_γ8/C2 (red) complex. Diameter differences at the Q/R site are mainly caused by conformational variations at R586 side chain among these three models. **c**, Pore dimensions measured between Cα of GluA1 Q582 and GluA2 R586 in resting state A1/2_γ8 (left), A1/2_γ8/C2 (middle) and active state A1/2_γ8/C2 (right). Upon receptor activation, the distance between GluA2 R586 is increased by ~1 Å in A1/2_γ8/C2. **d**, Distance measured between Cα of GluA1 C585 and GluA2 C589 at A1/2_γ8 resting state (left), A1/2_γ8/C2 resting (middle) and active (right) state. The corresponding EM densities are shown as mesh. Upon receptor activation, the distance between GluA2 C589 also increased by ~1.5 Å in A1/2_γ8/C2. All diameter labels are measured **in** Å. **e**, Charge distribution maps of the intracellular face of A1/2_γ8/C2 (red: -5 $k_BT/e$, blue: 5 $k_BT/e$) in the resting (top) and active (bottom) state indicate a dilation of the pore entrance in the direction of GluA2, but not GluA1 during receptor activation.

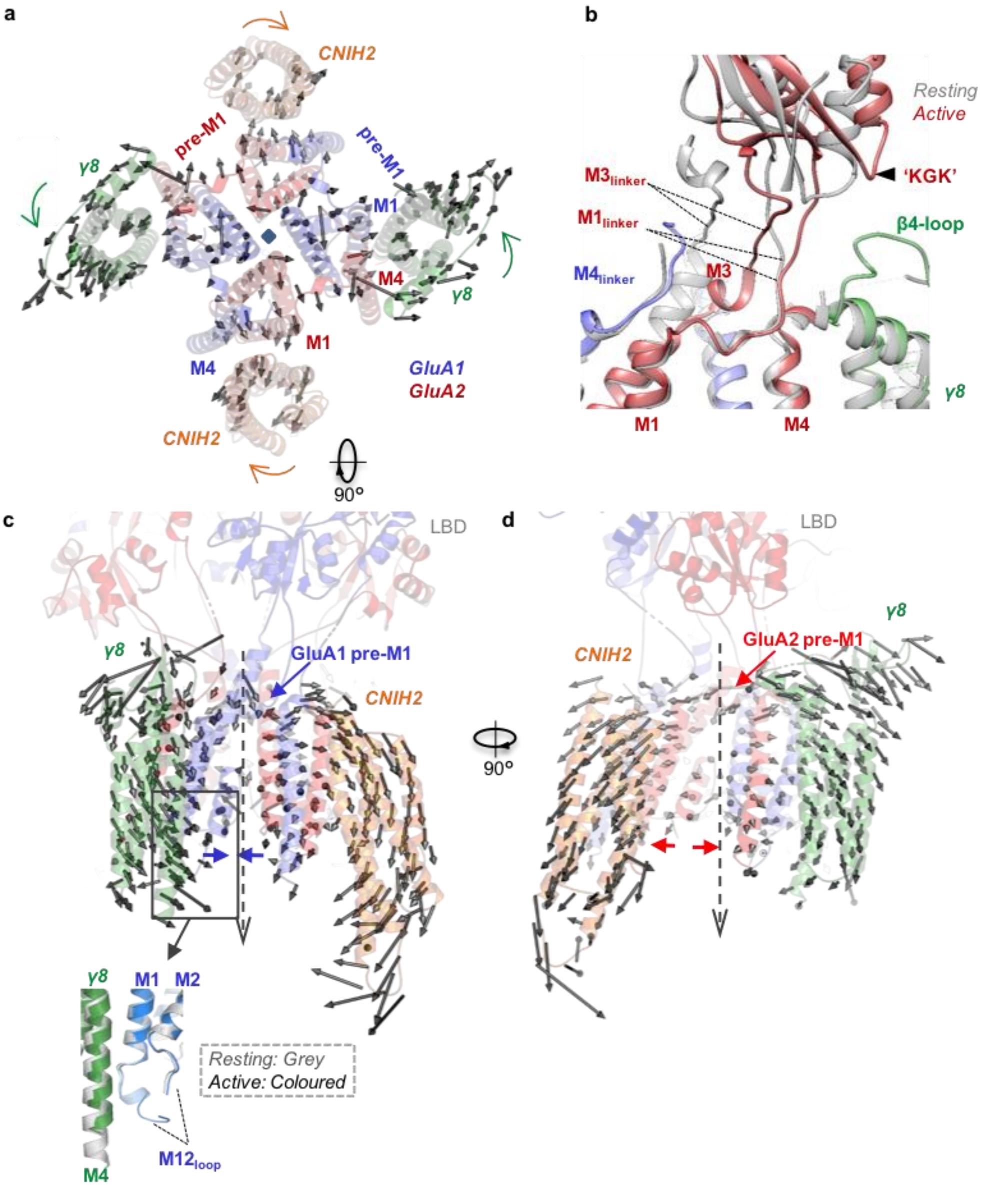


**Extended Data Figure 8 | Conformational changes of A1/2_γ8/C2 during receptor activation. a**, Top view superposition of A1/2_γ8/C2 in resting (grey) and active (red) states shows dilation of receptor and rotation of γ8 and C2 during activation. **b**, Superposition of A1/2_γ8/C2 in resting (grey) and active (coloured) states shows the conformational change of GluA2 M1 and M3 linkers as well as the LBD region upon receptor activation. The GluA2 M3 linker moves towards M1 linker, while the latter approaches γ8 'acidic' β4 loop. The LBD 'KGK' motif also moves towards the γ8 'acidic' loop. **c, d**, Conformational change of γ8 and C2 during receptor activation. The translation of the Cα atoms from the resting to active state is indicated as arrows for every second residue. Arrows indicate the direction and distance of helical movements. Auxilliary subunits come together on the GluA1 pre-M1 side (**c**), but are separated on the GluA2 pre-M1 side (**d**). Zoomed panel (**c**) indicates a contact between the γ8 M4 helix and the base of the GluA1 M1/2 cytoplasmic loop formed during receptor activation.

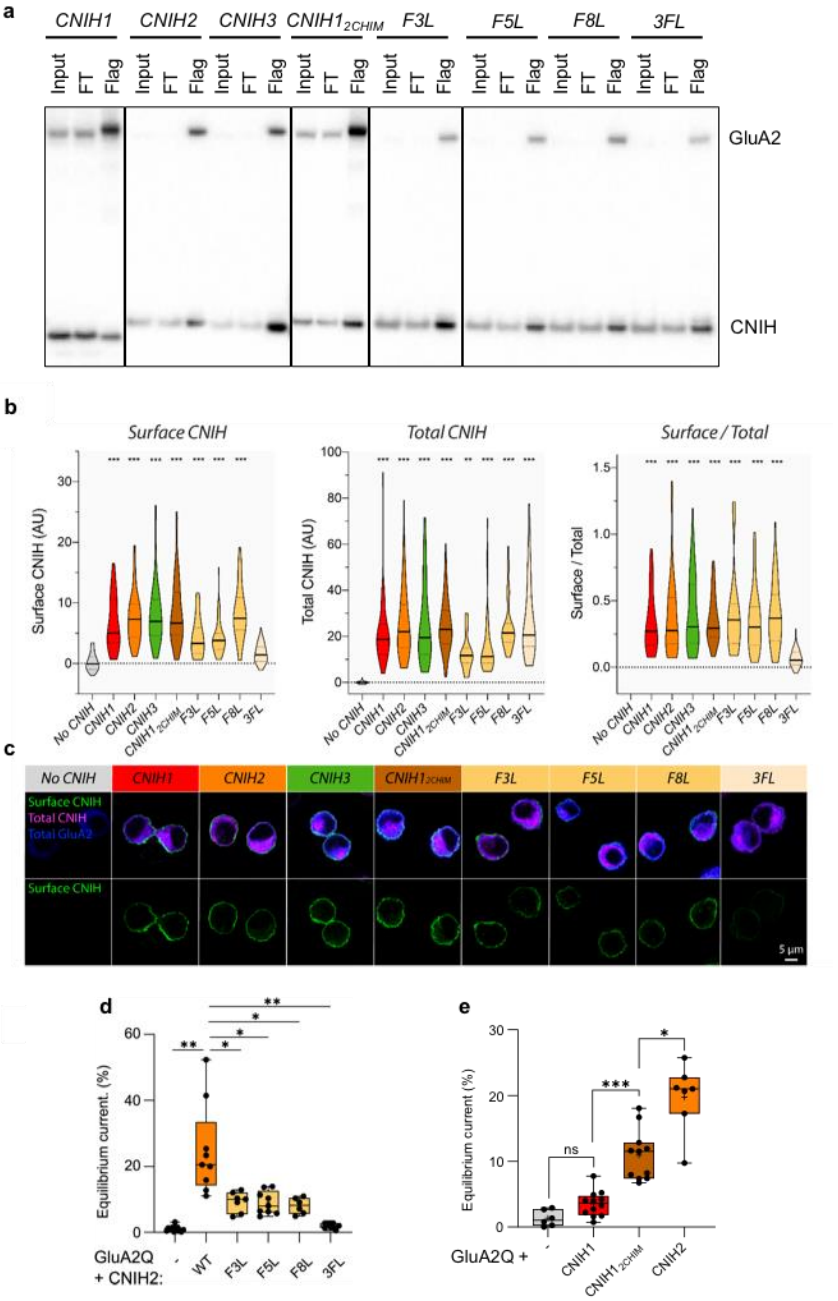


**Extended Data Figure 9 | Flag IP, immunostaining, and electrophysiology of CNIH homologues and CNIH2 mutants in complex with GluA1 or GluA2 homomers. a**, Flag IP of CNIH homologues and CNIH2 mutants in complex with GluA2 homomers. $CNIH1_{2CHIM}$, CNIH 1 and CNIH2 chimera with a fragment of CNIH2 (51-RERLKNIERICCLLRK-66) inserted into CNIH1 between P50 and L51; F3L, CNIH2 F3L; F5L, CNIH2 F5L; F8L, CNIH2 F8L; CNIH2 3FL, mutate all three phenylalanine 3, 5, 8 in CNIH2 to leucine; FT, flow through. **b**, Surface CNIH fluorescence (left), Total CNIH fluorescence (middle) and Surface/Total ratio (right) for CNIH homologues and CNIH2 mutants in complex with GluA2 (Surface CNIH (AU) – No CNIH: 0.21 ± 0.15, n=80; CNIH1: 6.60 ± 0.63, n=46; CNIH2: 7.54 ± 0.56, n=55; CNIH3: 8.07 ± 0.64, n=61; $CNIH1_{2CHIM}$: 8.56 ± 0.69, n=61; F3L: 4.42 ± 0.80, n=17; F5L: 4.25 ± 0.40, n=50; F8L: 8.36 ± 0.75, n=34; 3FL: 1.67 ± 0.22, n=50; Welch's ANOVA with Dunnett's multiple comparisons test: W(8,299.3) = 39.2, p<0.0001. Total CNIH (AU) – No CNIH: 0.02 ± 0.06, n=80; CNIH1: 21.5 ± 2.03, n=46; CNIH2: 25.8 ± 2.08, n=55; CNIH3: 25.2 ± 2.19, n=61; $CNIH1_{2CHIM}$: 25.0 ± 1.49, n=61; F3L: 12.7 ± 1.85, n=17; F5L: 16.2 ± 1.97, n=50; F8L: 24.6 ± 2.00, n=34; 3FL: 27.3 ± 2.42, n=50; Welch's ANOVA with Dunnett's multiple comparisons test: W(8,128.7) = 128.3, p<0.0001. Surface/Total – CNIH1: 0.34 ± 0.03, n=46; CNIH2: 0.39 ± 0.04, n=55; CNIH3: 0.42 ± 0.04, n=61; $CNIH1_{2CHIM}$: 0.36 ± 0.02, n=61; F3L: 0.41 ± 0.08, n=17; F5L: 0.35 ± 0.03, n=50; F8L: 0.39 ± 0.04, n=34; 3FL: 0.07 ± 0.01, n=50; Welch's ANOVA with Dunnett's multiple comparisons test: W(7,119.3) = 49.5). Homologues CNIH1, CNIH2 and CNIH3 show robust surface expression. CNIH2 mutants F3L, F5L and F8L, as well as the $CNIH1_{2CHIM}$ chimera, also traffic to the cell surface, whereas 3FL does not. F3L and F5L CNIH2 mutants show decreased total and, consequently, surface expression levels; to ensure the AMPARs in our electrophysiology experiments were still saturated with CNIHs we used a 1:2 AMPAR:CNIH co-transfection ratio. Increasing this ratio further to 1:4 for F3L & F5L did not affect the gating properties, suggesting that the observed change in AMPAR modulation by these mutants is not caused by their lower (surface) expression. **c**, Representative images showing surface CNIH (green), total CNIH (magenta) and total GluA2 (blue). **d**, Equilibrium current (**Fig. 3a** data set): (% peak) – GluA2 alone: 1.03 ± 0.19, n=15; CNIH2 WT: 24.72, ± 4.55, n=9; F3L: 9.25 ± 1.16, n=7; F5L: 8.96 ± 1.16, n=9; F8L: 8.05 ± 1.00, n=6; 3FL: 2.01 ± 0.28, n=9; Welch's ANOVA with Dunnett's multiple comparisons test: W(5,17.48) = 27.95, p<0.0001). **e**, Equilibrium current (**Fig. 3c** data set): GluA2 alone: 1.33 ± 0.50, n=6; CNIH1: 3.52 ± 0.56, n=12; $CNIH1_{2chimera}$: 10.93 ± 1.16, n=11; CNIH2: 19.77 ± 1.93, n=7; Welch's ANOVA with Dunnett's multiple comparisons test, W(3,15.36) = 40.08, p<0.0001).

**Extended Data Table 1.** Cryo-EM data collection, refinement and validation statistics.

| | A1/2_γ8/C2 Resting state | | A1/2_γ8 Resting state | A1/2_γ8/C2 Active state |
|---|---|---|---|---|
| | LBD-TMD | NTD | LBD-TMD | LBD-TMD |
| **Data collection and processing** | | | | |
| Microscope | | FEI Titan Krios | | FEI Titan Krios |
| Detector | | K3 + GIF | | K3 + GIF |
| Magnification | | 81000X | | 81000X |
| Voltage (kV) | | 300 | | 300 |
| Electron exposure (e-/Å2) | | 50 | | 50 |
| Defocus range (μm) | | -1.2 to -2.4 | | -1.2 to -2.4 |
| Pixel size | | 1.1 | | 1.1 |
| Symmetry imposed | | C2 | | C2 |
| Micrographs | | 10462 | | 8246 |
| Map resolution (Å) | 3.2 | 3.5 | 3.6 | 3.7 |
| FSC threshold | 0.143 | 0.143 | 0.143 | 0.143 |
| **Refinement** | | | | |
| Initial model used (PDB) | 6QKC | 6QKC | 6QKZ | 6QKC |
| Model resolution (Å) | 3.1 | 3.4 | 3.5 | 3.5 |
| FSC threshold | 0.5 | 0.5 | 0.5 | 0.5 |
| Map sharpening B factor (Å$^2$) | -113 | -142 | -115 | -122 |
| Model composition | | | | |
| Non-hydrogen atoms | 18022 | 11312 | 13648 | 17272 |
| Protein residues | 2322 | 1480 | 1922 | 2304 |
| Ligands | NBQX: 4 | NAG: 18, BMA: 2 | NBQX: 4 | CTZ: 4 |
| Lipids | 44 | 0 | 6 | 26 |
| B factors (Å2) | | | | |
| Protein | 66.42 | 13.70 | 80.43 | 50.86 |
| Ligand | 60.30 | 26.94 | 70.74 | 39.92 |
| R.m.s. deviations | | | | |
| Bond lengths (Å) | 0.005 | 0.008 | 0.005 | 0.009 |
| Bond angles | 0.600 | 0.578 | 0.605 | 0.847 |
| Validation | | | | |
| Molprobity score | 1.31 | 1.62 | 1.51 | 1.66 |
| Clashscore | 3.30 | 4.37 | 5.19 | 6.45 |
| Poor rotamers (%) | 0 | 0 | 0 | 0 |
| Ranachandran plot | | | | |
| Favored (%) | 96.88 | 94.01 | 96.49 | 95.66 |
| Allowed (%) | 3.12 | 5.99 | 3.51 | 4.34 |
| Disallowed (%) | 0 | 0 | 0 | 0 |
